# The causes of signed linkage disequilibrium within genomic datasets

**DOI:** 10.64898/2026.04.17.719204

**Authors:** Roman Stetsenko, Claire Mérot, Sylvain Glémin, Denis Roze

## Abstract

The signed linkage disequilibrium (LD) among selected mutations plays an important role in theoretical models of the evolution of sex and recombination. Several recent studies have quantified LD among mutations in genomic datasets, often reporting positive LD, particularly among mutations presumed to be less deleterious, such as synonymous variants. In this article, we investigate two potential sources of this positive LD: the focus on rare alleles, as adopted in several previous studies, and errors arising in the mapping of short-read sequences onto a reference genome. Using coalescent simulations, we extend previous theoretical results of the effect of focusing on rare alleles, and show that derived neutral alleles present at similar frequencies tend to be in positive LD, while alleles present at different frequencies tend to be in negative LD. Reanalyzing datasets from *Capsella grandiflora* and *Drosophila melanogaster*, we show that LD among synonymous derived alleles vanishes in the absence of any conditioning on frequency, while LD between mutations categorized as potentially deleterious by the SIFT4G program stays positive. However, we show that in both cases, this positive LD may be at least partly caused by the potential mismapping of a small fraction of sequences in some individuals, which could be a consequence of structural variants that are absent from the reference genome. Overall, these results show that average signed LD among mutations can be strongly affected by technical artifacts even if these concern only a minority of variants. Finally, we discuss other possible sources of positive LD among deleterious mutations.

## Introduction

Deleterious mutations may be abundant within natural populations (Charlesworth, 2015; Lynch et al., 1999), and are thought to be involved in various evolutionary processes such as the evolution of aging (Flatt and Partridge, 2018), mating systems (Charlesworth, 2006; Lande and Schemske, 1985), sex and recombination (Agrawal, 2006; Otto, 2009; Otto and Lenormand, 2002). The linkage disequilibrium between deleterious alleles at different loci (measuring their degree of association within genomes) can play an important role in several of these processes, the evolution of sex and recombination in particular (Felsenstein, 1974; Otto, 2021; Otto and Lenormand, 2002). Given two alleles *a* and *b* at two polymorphic loci, the linkage disequilibrium (LD) between those alleles is classically defined as: *D_ab_* = *p_ab_* −*p_a_p_b_*, where *p_ab_* is the frequency of haplotypes *ab* in the population, and *p_a_*, *p_b_* the frequencies of alleles *a* and *b* at each locus (Lewontin and Kojima, 1960; Weir, 1996). *D_ab_* thus equals zero when allele *a* is not more (or less) associated with allele *b* than expected based on the frequency of allele *b* in the population, while *D_ab_* is positive (negative) when the frequency of the *ab* haplotype is higher (lower) than expected based on allele frequencies. In other words, a positive *D_ab_* means that *a* and *b* are more often found together on the same haplotype. If *a* and *b* are neutral, the expected value of *D_ab_* is zero at mutation-recombination-drift equilibrium; however, random drift generates a variance in *D_ab_*, so that the expected squared LD (denoted ^+^*D_ab_*^2,^ hereafter) is positive. In a randomly mating population, ^+^*D_ab_*^2,^ is approximately (10 + *ρ*) ⟨*p_a_q_a_p_b_q_b_*⟩ */* [(2 + *ρ*) (11 + *ρ*)] at equilibrium where *ρ* = 4*N_e_ r_ab_*, *N_e_* is the effective population size, *r_ab_*the recombination rate between the two loci, and *q_i_* = 1 − *p_i_* (Hill and Weir, 1988; McVean, 2002; Ohta and Kimura, 1971). This quantity tends to ⟨*p_a_q_a_p_b_q_b_*⟩ */ρ* when *ρ* is large. Squared LD is often measured in genomic studies in order to assess the range of genetic distances over which genetic associations (regardless of their sign) are maintained, informative on a broad range of evolutionary processes (e.g., the effect of recombination, drift, selection, population structure, the mating system; reviewed by Slatkin, 2008). For example, genetic associations are typically maintained up to approximately 100 kb in humans (Reich et al., 2002) and 1 kb in *Drosophila melanogaster* (MacKay et al., 2012).

If alleles *a* and *b* are deleterious, selection is more efficient when LD is positive since the variance in fitness in the population is higher than when LD is negative. As a consequence, random drift and selection tend to generate negative LD (on average) between deleterious alleles in finite haploid populations (selective interference or the Hill-Robertson effect: Felsenstein, 1974; Hill and Robertson, 1966; Otto, 2021). Indeed, situations with negative LD tend to persist longer than situations with positive LD, due to the lower variance in fitness when LD is negative. Epistasis between selected loci (when measured on a multiplicative scale) is another possible source of LD, generating LD of the same sign as epistasis (Felsenstein, 1965). Positive epistasis means that the deleterious alleles *a* and *b* tend to compensate each other when combined, leading to a higher frequency of the *ab* haplotype than expected based on the frequencies of *a* and *b* (*D_ab_ >* 0), while negative epistasis means that the deleterious effects of *a* and *b* tend to reinforce each other when combined, leading to a lower frequency of the *ab* haplotype (*D_ab_ <* 0). In the presence of negative LD, recombination tends to increase the variance in fitness among offspring, thereby increasing the efficiency of natural selection. However, recombination may also decrease the mean fitness of offspring, particularly when epistatic interactions are present (e.g., Agrawal, 2006; Otto and Lenormand, 2002).

Due to its central role in various evolutionary processes, several recent studies have tried to assess the overall sign and magnitude of LD (*D_ab_*) between putatively deleterious alleles segregating within populations. These studies typically contrast different types of mutations: synonymous, missense, and loss-of-function (LoF) mutations (Garcia and Lohmueller, 2021; Ragsdale, 2022; Sandler et al., 2021; Sohail et al., 2017; Stolyarova et al., 2022). Sohail et al. (2017) estimated the genome-wide LD (corresponding to the sum of LD among all pairs of mutations) for each class of mutation from the under- or overdispersion of the distribution of the number of mutations per genome (given that this distribution should be approximately Poisson in the absence of LD), in *Drosophila melanogaster* and humans. They found that genome-wide LD is positive between synonymous (rare) mutations, less positive between missense mutations and negative between LoF (the presumably most deleterious) mutations, which they interpreted as a possible effect of negative epistasis between deleterious mutations. The *D. melanogaster* dataset was later reanalyzed by Sandler et al. (2021), who also analyzed an additional dataset from a single population of the outcrossing plant *Capsella grandiflora*. They found genome-wide positive LD for all classes of (rare) mutations except LoF mutations in *D. melanogaster* which displayed negative LD; however, the latter was found to be non-significant. Additionally, they computed LD for different classes of distances (in bp) between pairs of loci, showing that at short distances, LD between LoF mutations is more positive than between synonymous mutations. Sandler et al. (2021) hypothesized that this pattern may be explained by positive intragenic epistasis, additional LoF mutations having little effect once a gene is already disrupted by a first LoF mutation. Garcia and Lohmueller (2021) found that non-synonymous rare variants exhibit more negative LD than synonymous rare variants in human populations, and interpreted this result as the consequence of selective interference (Hill-Robertson effect) among deleterious alleles. However, the measure of LD used (*D^′^*) makes it difficult to compare their results with those obtained in previous studies, and generates a bias towards finding negative LD (Johri and Charlesworth, 2025). Ragsdale (2022) used genomic data from 15 human populations to measure LD within protein-coding genes, and found positive LD between derived synonymous and missense mutations, but slightly negative LD between LoF mutations. Finally, Stolyarova et al. (2022) found higher positive LD among nonsynonymous than among synonymous variants within populations of the hypervariable fungus *Schizophyllum commune*, which was again interpreted as evidence for positive epistasis among segregating deleterious mutations.

Importantly, in several of these studies LD was measured among rare variants, since deleterious alleles are expected to stay at low frequency within populations (Garcia and Lohmueller, 2021; Sandler et al., 2021; Sohail et al., 2017). However, a theoretical study by Good (2022) showed that such a conditioning on allele frequencies generates a bias towards positive LD, which is stronger in the case of neutral alleles than in the case of deleterious alleles (see also Johri and Charlesworth, 2025). Additionally, Sandler et al. (2021) used a simulation model to show that past admixture may also generate positive LD among rare neutral variants, the effect of admixture being much reduced in the case of deleterious variants. These results may thus explain the positive LD observed among synonymous variants, and show that observing less positive LD among missense than among synonymous rare variants does not necessarily imply the existence of a selective force generating negative LD among deleterious alleles within populations (Sandler et al., 2021). However, they do not explain situations in which LD among closely linked missense variants is more positive than among synonymous variants (Ragsdale, 2022; Sandler et al., 2021; Stolyarova et al., 2022).

In this article, we explore and discuss possible sources of positive LD between variants in genomic datasets. Using coalescent simulations, we show that positive LD is expected (on average) among mutations present at similar frequencies within a population or a sample (not only rare mutations). By re-analyzing the sequence data from *Capsella grandiflora* and *Drosophila melanogaster* used by Sandler et al. (2021), we show that the positive LD among derived neutral variants disappears in the absence of any conditioning on allele frequency. While positive LD is still observed among putatively deleterious variants in both datasets, we show that this positive LD may at least partly be caused by mapping errors, that inevitably occur when using short-read sequencing data (e.g., Igolkina et al., 2025). In the *C. grandiflora* dataset, these may result from structural variants segregating within the population and generating pseudo-heterozygosity, as found by Jaegle et al. (2023) on data from the 1001 Arabidopsis Genome Project. In the *D. melanogaster* dataset (obtained from haploid embryos and in which pseudo-heterozygous sites have been filtered out), we show that a substantial proportion of the positive LD observed among putatively deleterious variants is caused by a family of duplicated genes present on the X chromosome, which may also have caused mapping errors. Finally, we discuss other possible sources of positive LD among selected mutations.

## Results

### Neutral alleles at similar frequencies tend to be in positive LD

Good (2022) showed that in a finite, panmictic population, the linkage disequilibrium between rare neutral mutations is positive on average. Our coalescent simulations show that this result can be extended to the case of neutral variants segregating at similar frequencies in a population (not only rare variants). As shown by Figure 1, the average LD among derived alleles is zero when computed over all pairs of segregating alleles in the sample (no conditioning on frequency: grey dots), while it is positive when computed over pairs of loci at which derived alleles are present at similar frequencies (both rare or both frequent). One can note that the average LD between frequent derived alleles is higher than between rare derived alleles (compare Figures 1A and 1B). This result can be understood from the fact that mutations present at similar frequencies are more likely to have arisen in the same region of the coalescent tree, and thus to be in positive LD. This is perhaps most easily seen in the case of mutations present in *n* − 1 sequences (where *n* is the sample size): in this case, assuming no recombination, both mutations necessarily arose in the branch between the last common ancestor of these *n* − 1 sequences and the coalescence event between this branch and the ancestral branch of the other sequence, so that the two mutations are always present in the same sequences (Figure S1). This bias towards positive LD is less strong in the case of mutations present at lower frequencies, but is still present. This effect of frequency may also be seen by considering the extreme cases where only *AB* and *ab* haplotypes are present (maximum positive LD) or where only *Ab* and *aB* haplotypes are present (maximum negative LD): in the first case, *a* and *b* are necessarily at the same frequency, while in the second case, *a* is frequent if *b* is rare (and vice versa). Figure S2 shows that derived alleles present at different frequencies are more often in negative than in positive LD (although LD may be positive on average between alleles present in the highest frequency class and alleles present at lower frequencies, as shown by Figure S2F). The positive LD between variants present at similar frequencies and the negative LD between variants present at different frequencies cancel out when averaging over all variants, so that the average LD is zero in the absence of any conditioning on frequency. When LD is scaled by products of allele frequencies for each pair of segregating variants, the resulting average is not necessarily zero, however: indeed, Figure S3 shows that the ratio *D_ab_/p_a_q_a_p_b_q_b_* is positive when averaged over all pairs of variants (by contrast, the average of 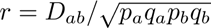 remains close to zero, as shown by Figure S4).

**Figure 1:**
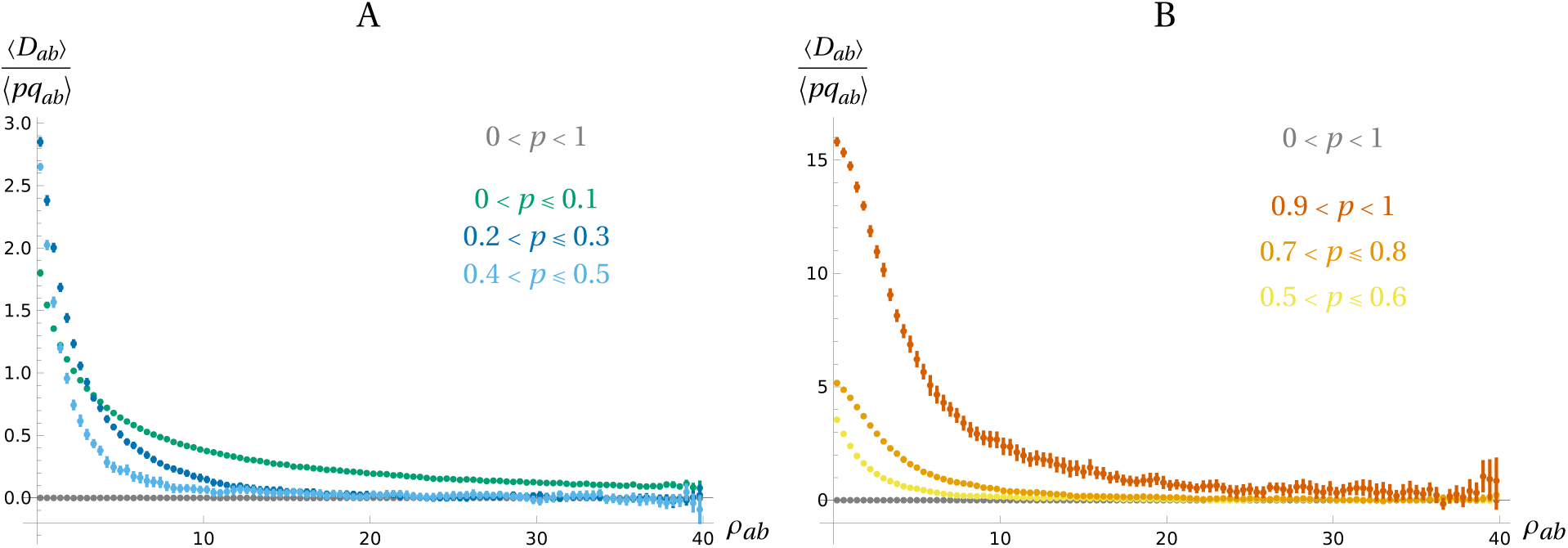
Average LD 〈*D_ab_*〉 between derived neutral mutations (scaled by the average product of genetic diversities 〈*pq_ab_*〉 = 〈*p_a_q_a_p_b_q_b_*〉) in the coalescent simulations, as a function of the recombination rate *ρ* = 4*N_e_r_ab_*. In color: LD is computed over pairs of loci at which the derived allele segregates in a given frequency range; In grey: LD is computed between all derived alleles, irrespective of their frequency. Errors bars correspond to 95% confidence intervals (see Methods section). Parameters values: *θ* = 4*N_e_u* = 100 (mutation rate); *n* = 400 (sample size).

As shown by Figure S5, polarizing LD based on minor allele frequency (MAF) — that is, computing LD among alleles whose frequency is below a given threshold, independently of the ancestral or derived status of alleles — generates the same bias towards positive values, which is more important when the threshold frequency is lower. Results on the average squared LD ^+^*D_ab_*^2,^ are shown on Figures S6 – S7: in the absence of any conditioning on frequency, ^+^*D_ab_*^2,^ is well predicted by (10 + *ρ*) ⟨*p_a_q_a_p_b_q_b_*⟩ */* [(2 + *ρ*) (11 + *ρ*)] (Figure S6A; Hill and Weir, 1988; McVean, 2002; Ohta and Kimura, 1971). However, the squared LD between variants in the same frequency class is either lower (for rare variants) or higher (for frequency classes higher than 0.2; Figure S6B). When LD is polarized based on minor allele frequency, the squared LD is lower than the theoretical prediction (Figure S7).

Figures S8 and S9 show that introducing a given rate *µ* of misspecification of ancestral/derived state does not affect much the previous results, the average LD staying at zero in the absence of conditioning on allele frequencies (Figure S8A), and positive when measured only among ‘derived’ alleles present at similar frequencies (Figures S8B and S8C). Misspecification decreases the average LD among frequent ‘derived’ alleles (Figure S8D), since this class now contains LD measures between rare (and truly) derived alleles. For the same reason, misspecification decreases the squared LD among frequent ‘derived’ alleles (Figure S9D) whereas it does not effect much the results in the absence of conditioning on allele frequency (Figure S9A) and when measured only among ‘derived’ alleles present at intermediate frequencies (Figures S9B and S9C).

Our simulation results therefore predict that LD between neutral derived variants should be zero on average, in the absence of any conditioning on allele frequency, while LD between neutral variants present at similar frequencies should be positive on average. Using genome-wide SNP data from 182 individuals of a single population of the obligatory outcrossing plant *Capsella grandiflora* and 190 haploid embryos of *Drosophila melanogaster*, we measured LD between putatively neutral (SIFT score = 1) derived variants (see Methods section). In the absence of any conditioning on allele frequency we found, as expected, null LD on average between neutral derived variants for all distance classes, in both species (Figure 2). Similarly to our simulation results, we found that in both datasets, LD between variants present at similar frequencies is positive on average (Figure 2 and Figure S10). In particular, we found similar qualitative patterns than in our simulations when LD was measured between derived variants in the same frequency class (Figure 2) or with different MAF thresholds (Figure S10). The average squared LD also showed similar patterns as in our simulations when conditioning on the frequency of derived variants (Figure S11 and Figure S12).

**Figure 2:**
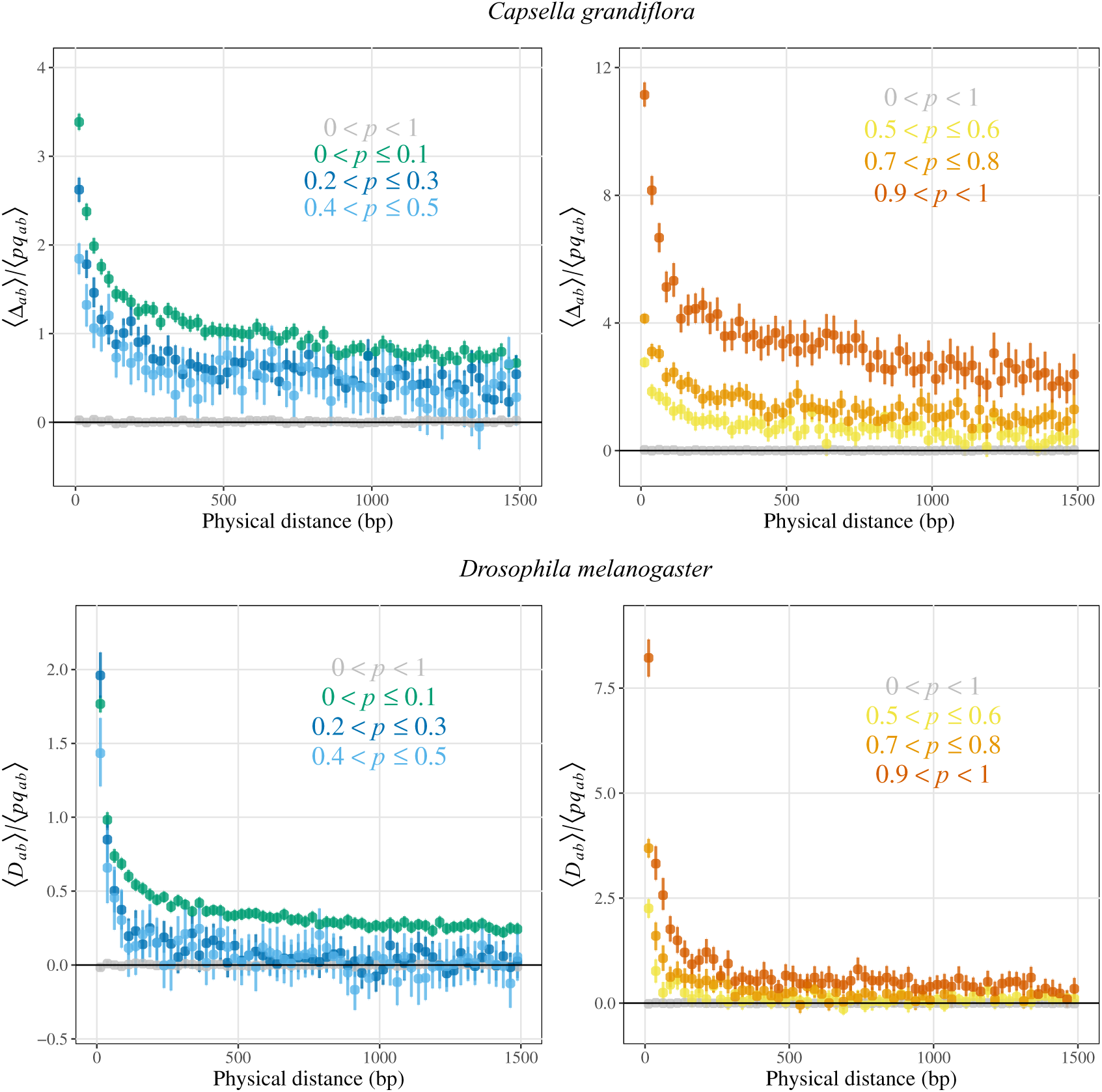
Average composite linkage disequilibrium Δ*_ab_* or linkage disequilibrium *D_ab_* (scaled by the average product of allele frequencies) between neutral derived variants according to physical distance between variants, in *C. grandiflora* and *D. melanogaster*. In color: LD is computed over pairs of loci at which the derived allele segregates in a given frequency range (same as in Figure 1; In grey: LD is computed between all neutral derived variants, irrespective of their frequency.

### Positive LD between putatively deleterious mutations in *C. grandiflora* and *D. melanogaster*

As explained in the Methods section, alleles at polymorphic sites were categorized as ‘neutral’, ‘mildly deleterious’ or ‘deleterious’ based on their SIFT score (Vaser et al., 2016). The site frequency spectra (SFS) of derived mutations reflected their SIFT score category in both species as deleterious mutations had an SFS shifted towards rarer frequencies compared to mildly deleterious mutations and furthermore compared to neutral mutations (Figure S13).

As shown by Figure 3, the average LD between deleterious alleles and between mildly deleterious alleles is positive between pairs of variants located at small physical distance, in both *C. grandiflora* and *D. melanogaster*. In *D. melanogaster*, the average LD drops to zero as the distance between sites increases (and similarly in the case of mildly deleterious mutations in *C. grandiflora*), while LD between deleterious mutations stays positive at least up to 5-10kb in *C. grandiflora* (Figure S14). This positive LD is not caused by any conditioning on the frequency of variants, since averages are computed over all deleterious and mildly deleterious variants (without any restriction based on frequency). Restricting the analyses to derived deleterious and mildly deleterious variants (defining an allele as ancestral if it is present in the outgroup *N. paniculata* or *D. yakuba*) does not change significantly these patterns of LD, as most deleterious variants are derived (results not shown). The LD pattern shown in Figure 3 is mostly driven by associations between mutations present in the same gene, since at such short distances, a large majority (95% for *C. grandiflora* and 94% for *D. melanogaster*) of pairs of mutations are found within the same gene (Figure S15). Due to the small number of pairs of mutations located in different genes (particularly at short distances), it is not possible to conclude on a difference in LD patterns within vs. between genes (a similar result was found by Sandler et al., 2021 and Ragsdale, 2022).

**Figure 3:**
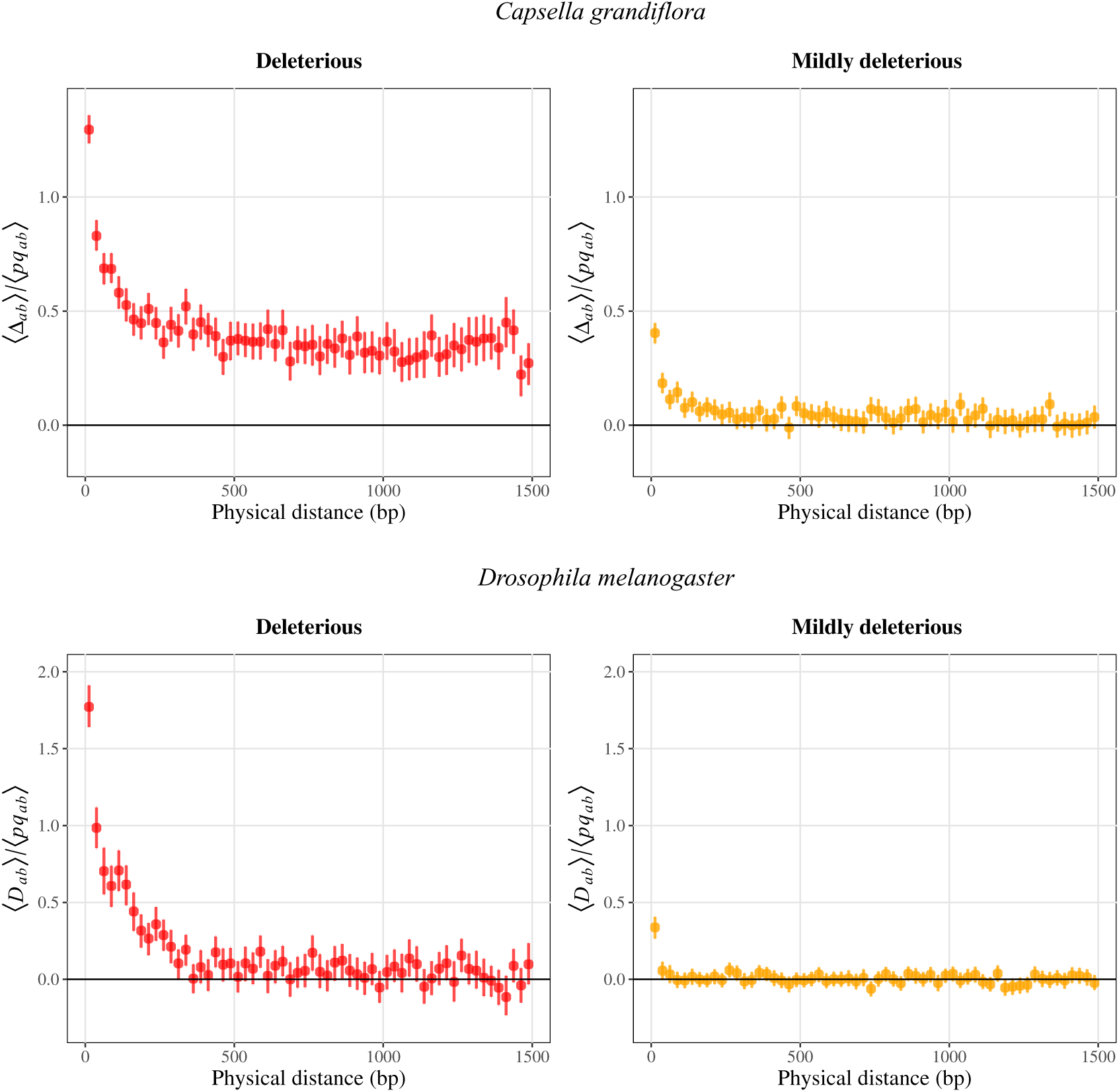
Average composite linkage disequilibrium Δ*_ab_* or linkage disequilibrium *D_ab_* (scaled by the average product of allele frequencies) between deleterious or mildly deleterious variants according to physical distance in *C. grandiflora* and *D. melanogaster*.

We also used the *C. grandiflora* and *D. melanogaster* datasets to test theoretical predictions on moments of allele frequencies and linkage disequilibrium. In particular, Roze (2021) showed that the quantity ⟨*p_b_D_ab_*⟩ should be negative when allele *b* is neutral while allele *a* is deleterious: indeed, when allele *b* tends to be associated with the deleterious allele *a* (that is, when *D_ab_ >* 0), the frequency of *b* should decrease. By contrast, ⟨*p_a_D_ab_*⟩ should be zero on average, since being associated with a particular neutral allele should not affect the frequency of a deleterious allele. Our results tend to confirm these predictions, as we observe a negative correlation between the frequency of neutral alleles and their LD with deleterious alleles at short genetic distances in *C. grandiflora* and very short distances (<100 bp) in *D. melanogaster* (Figure S16).

### Positive LD between putatively deleterious mutations may at least partly be due to mapping errors

As illustrated by Figure S17, mapping errors are a possible source of pseudo-heterozygosity that may give rise to spurious LD among mutations. Such mapping errors may be caused by polymorphic duplications present in some of the sequenced individuals but not in the reference genome, and incorrectly mapped onto the original sequence (see Figure S17; Jaegle et al., 2023). The consequences of pseudo-heterozygosity may be more important in the case of deleterious than in the case of neutral mutations, since deleterious mutations are less abundant within genomes, and a significant proportion of these mutations may be wrongly categorized as deleterious due to such mapping errors. In the case of polymorphic duplications, for example, mutations in a duplicated gene may not be deleterious if a functional gene copy is maintained elsewhere in the genome. Pseudo-heterozygosity is not an issue in the *D. melanogaster* dataset as sequences were obtained from haploid embryos (see Methods). By contrast, pseudo-heterozygosity may potentially contribute to the positive LD observed among deleterious mutations in *C. grandiflora*.

As explained in the Methods, we developed an *R* program to identify potential blocks of pseudo-heterozygosity (containing multiple variants that are present only in the heterozygous state, in the same individuals), that we ran on the SNPs called in *C. grandiflora* individuals. In *C. grandiflora*, we identified a large number of potential blocks of pseudo-heterozygosity (92,817). We restricted our analyses to 155 blocks comprising at least 5 deleterious mutations and showing a significant increase (at least a 1.3 fold) in coverage in individuals heterozygous for the block compared to homozygous individuals (as expected in the case of pseudo-heterozygosity caused by mapping errors). Those blocks are on average 1.7kb long (Figure S18A), carrying 18.3 SNPs (Figure S18B) and 7.9 deleterious mutations (Figure S18C). Heterozygous individuals for these blocks are in frequency 0.19 on average, with a frequency spectrum highly skewed towards low values (median frequency of 0.11; Figure S18D). A substantial proportion of these heterozygosity blocks are thus relatively frequent in the population, and the mutations categorized as deleterious by SIFT and present in these blocks may thus contribute significantly to the positive LD shown in Figure 3 (since loci where allele frequencies are closer to 0.5 make stronger contributions to average LD), while the fact that these mutations are present in relatively high frequency indicates that they are likely not deleterious. However, removing deleterious mutations present above a threshold frequency from the analysis would generate the same bias towards positive LD as shown in the case of neutral mutations above. Indeed, our “deleterious” mutations category necessarily includes at least some mutations that are neutral or nearly-neutral, and positive LD is expected (on average) among neutral mutations present below a threshold frequency (see Figures 1 and S5). When SNPs located in filtered heterozygosity blocks are excluded from LD computation, positive LD between deleterious mutations decreases, while positive LD between mildly deleterious is mostly unaffected (Figure 4 and S19). This suggests that at least part of the positive LD found between deleterious mutations in *C. grandiflora* may be caused by potential spurious deleterious mutations found in these pseudo-heterozygosity blocks.

**Figure 4:**
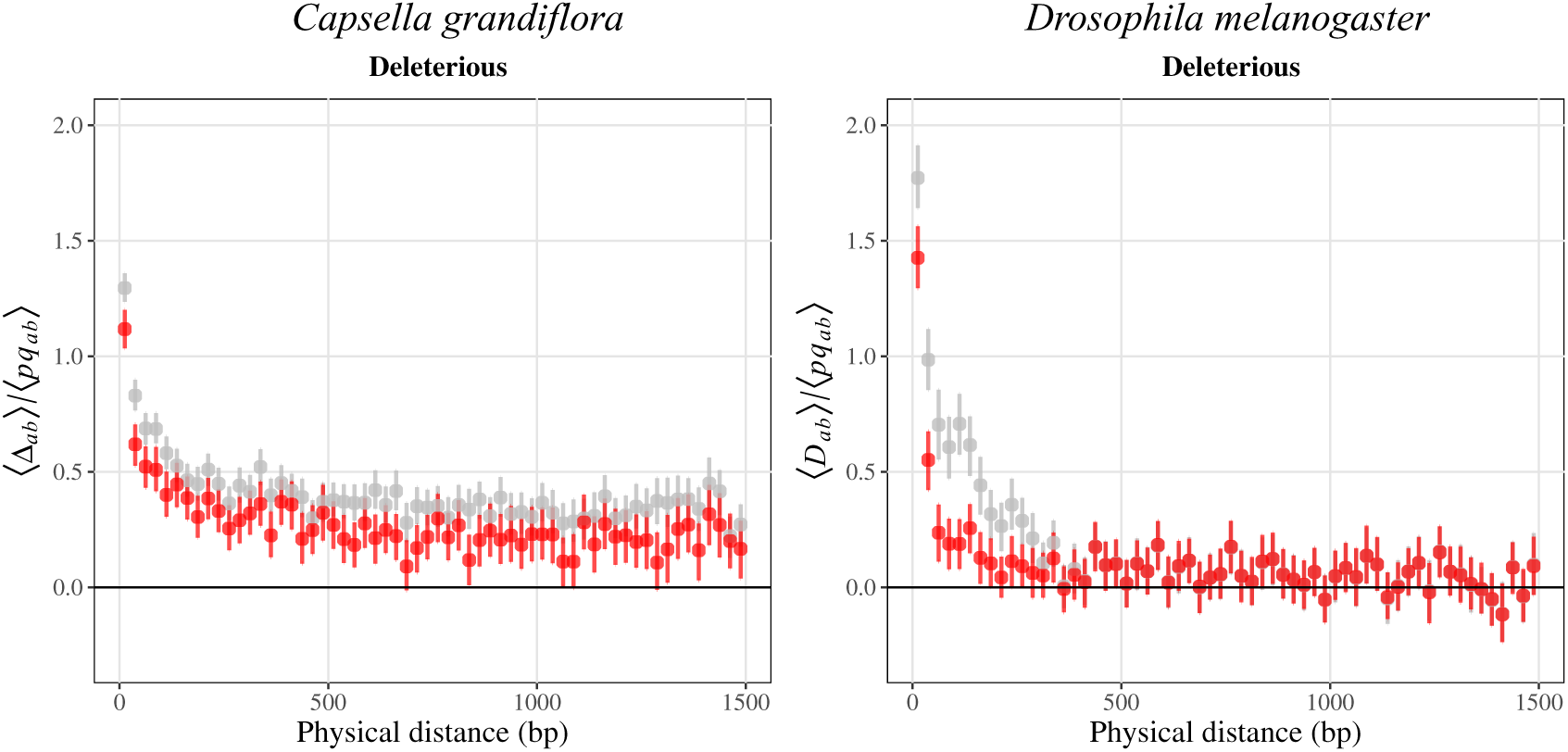
Left panel: Composite linkage disequilibrium Δ*_ab_* scaled by the product of allele frequencies between deleterious variants according to physical distance between sites, after removing SNPs located in heterozygosity blocks (coloured) or considering all SNPs (grey, as in Figure 3) in *C. grandiflora*. Right panel: Linkage disequilibrium *D_ab_* scaled by the product of allele frequencies between deleterious variants according to physical distance after removing 3 genes (CG33665, CG33666 and CG33669) on chromosome X (coloured) or considering all SNPs (grey, as in Figure 3) in *D. melanogaster*.

We also checked whether these heterozygosity blocks may correspond to potential structural variants called by programs using complementary information on paired-ends, split-reads and read-depth provided in short reads alignments. We found 51 blocks matching with SV calls, of which 16 matched with duplications calls (Table S2) — other matching SV calls included deletions and inversions, but we decided to focus on duplications since they are the most likely SV to generated pseudo-heterozygosity. Some of these validated heterozygosity blocks correspond to peaks of positive LD in the genome, although our method did not retrieve some of the highest peaks of positive LD (Figure 5). Examples of Samplot outputs showing SV calls, coverage pattern and the position of the heterozygosity block for heterozygous and homozygous individuals are shown in Figure S20, S21 and S22. These results thus suggest that the positive LD between deleterious mutations observed in *C. grandiflora* may be partly due to mapping errors caused by segregating duplications, that are absent from the reference genome of *C. rubella*.

**Figure 5:**
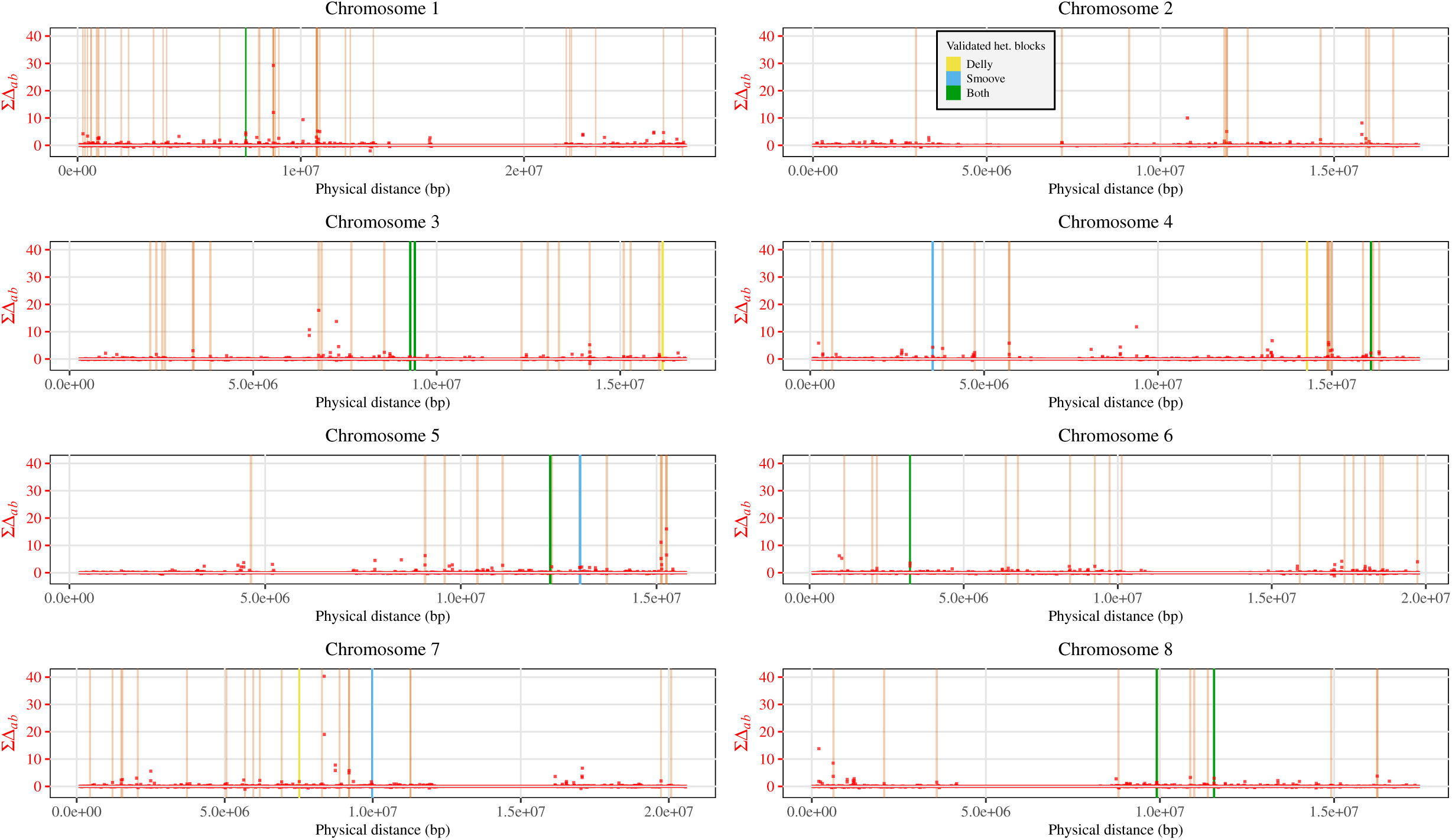
Sum of the composite linkage disequilibrium between pairs of SNPs ^Σ^ Δ*_ab_* (in 3kb windows; red points), along the genome of *C. grandiflora*. Heterozygosity blocks matching with duplication calls by Delly (yellow bars), Smoove (blue bars) or both (green bars) as well as the rest of heterozygosity blocks (orange bars) are also shown.

As mentioned above, pseudo-heterozygosity cannot contribute to the positive LD among deleterious mutations observed in the *D. melanogaster* dataset, as these sequences have been obtained from haploid embryos (Lack et al. 2015). However, detailed analysis showed that a substantial proportion of the observed positive LD is driven by 3 genes (CG33665, CC33666 and CG33669), which are part of a family of 6 paralogs (CG33664 – CG33669) present in the same region of the X chromosome. Indeed, Figure 4 and S19 show that removing these genes from the analysis substantially decreases the signal of positive LD, LD staying positive at short distances (<200bp). We observed that a variant sequence that differs from the sequences of the 6 paralogs at more that 25 nucleotides is present at intermediate frequencies in the dataset, being mapped on gene CG33665 in some individuals and on genes CC33666 or CC33669 in others, and generating the increase in positive LD shown in Figure 4 and S19. The exact cause of this pattern is difficult to assess: for example, it could be due to the presence of a diverged copy of one of the 6 paralogs in some individuals but absent from the reference genome, incorrectly mapped on one of the paralogs in these individuals. In any case, the fact that the variants causing the increase in positive LD are present at high frequency indicates that they are very unlikely to be deleterious.

## Discussion

Although linkage disequilibrium among deleterious mutations may play an important role in a number of evolutionary processes (the evolution of sex and recombination in particular; e.g., Otto, 2021; Otto and Lenormand, 2002), empirical efforts to quantify it from population genomic data remain relatively recent. A challenge in such estimations stems from the general difficulty in assessing whether a given variant is deleterious. Therefore, most authors have used a general approach that consists in comparing different broad classes of mutations, often contrasting synonymous, non-synonymous and loss-of-function (LoF) mutations (Garcia and Lohmueller, 2021; Sandler et al., 2021; Sohail et al., 2017; Stolyarova et al., 2022). Indeed, LoF mutations are expected to be on average more deleterious than non-synonymous mutations, while synonymous mutations should be approximately neutral (although a fraction of LoF and non-synonymous mutations may also be neutral, while some others may be advantageous or under balancing selection). Alternatively, several methods have been developed to predict whether an amino acid substitution is deleterious, such as the SIFT4G algorithm used in the present article and based on the degree of conservation of amino acids among homologous proteins (Vaser et al., 2016), or methods based on rates of substitutions of non-synonymous mutations within phylogenies (e.g., Davydov et al., 2010). Again, these methods are necessarily imperfect, and some of the predicted deleterious mutations may be neutral or under different forms of selection — however, Chen et al., 2022 showed that SIFT scores are consistent with fitness effects estimated using methods based on the site frequency spectrum. A second difficulty in estimating LD among deleterious mutations arises from possible biases or artifacts that may be difficult to detect. In the following, we discuss two such biases explored in this article, that tend to generate positive LD among mutations: the effect of focusing on rare alleles, and the effect of errors that inevitably occur when mapping short-read sequences onto a reference genome. Finally, we discuss possible biological sources of positive LD among deleterious alleles within natural populations.

A deleterious mutation is expected to stay at a low frequency, and several authors thus focused on rare variants to estimate LD among deleterious alleles (Garcia and Lohmueller, 2021; Sandler et al., 2021; Sohail et al., 2017). These studies typically found positive LD between rare variants, including synonymous mutations. One exception is the study by Garcia and Lohmueller (2021) who found negative LD (on average) between non-synonymous and between synonymous variants present at low frequency in human polymorphism data; however, the measure of LD used (*D^′^* as defined by Lewontin, 1964, which is scaled differently by products of allele frequencies for positive and negative LD) generates a bias towards finding negative LD (Johri and Charlesworth, 2025). Good (2022) provided analytical approximations for the average LD among rare alleles (his equations 37 and 40b), showing that it is positive (but less so for deleterious than for neutral variants), and vanishes to zero when the effect of drift on the frequency of deleterious alleles becomes negligible, in the absence of epistasis. Our coalescent simulation results show that in the case of neutral alleles, positive LD is expected (on average) between alleles present at similar frequencies (not only rare alleles), this effect being strongest in the case of high-frequency derived alleles. Conversely, LD between alleles present at different frequencies tends to be negative on average. As mentioned earlier, this effect can be understood from the fact that mutations present at similar frequencies are more likely to have arisen in the same region of the coalescent tree, and therefore to be in positive LD. One can note that the fact that deleterious mutations stay rare does not generate positive LD among them: indeed, deleterious mutations do not persist for long within populations, and thus tend to arise near the leaves of the coalescent tree. While previous studies on *Drosophila melanogaster* and *Capsella grandiflora* found positive LD between rare synonymous variants (Sandler et al., 2021; Sohail et al., 2017), our analyses based on the same datasets show that LD among neutral derived variants is zero on average in the absence of any conditioning on frequency. Furthermore, the average LD between mutations present at similar frequencies in these datasets shows similar patterns as in our coalescent simulations. Exploring to what extent patterns of LD as a function of allele frequencies could be used to infer past demographic or selective events would represent an interesting extension of this work. Overall, and in agreement with the previous works by Good (2022) and Johri and Charlesworth (2025), our results indicate that conditioning on allele frequency should be avoided when estimating LD among deleterious alleles segregating within populations. Indeed, although truly deleterious alleles should stay at low frequency, the variants categorized as deleterious will necessarily include a fraction of neutral and nearly neutral variants (as discussed above), and eliminating these variants only when present above a threshold frequency will generate a bias towards positive LD.

Our results also outline mapping errors as a possible source of spurious positive LD among mutations. Mapping errors have multiple possible causes, and may be difficult to detect. In particular, they can result from segregating duplications that are absent from the reference genome, generating patterns of pseudo-heterozygosity (Dallaire et al., 2023; Igolkina et al., 2025; Jaegle et al., 2023). In this case, nucleotides that differ between the original and duplicated sequence will appear as polymorphic due to the mismapping of the duplicated sequence onto the original one, with positive LD among the variant nucleotides present in the duplication. While this process may generate spurious positive LD between deleterious as well as between neutral mutations, its effect should be stronger in the case of deleterious mutations, for two reasons: (i) deleterious variants are expected to be less abundant within genomes than neutral ones, so that a stronger proportion of deleterious variants may result from such mapping errors, and (ii) deleterious variants should be present at low frequency, so that a few polymorphic duplications present at intermediate frequencies may have a strong effect on the average LD (since LD is stronger when loci are more polymorphic). This last point is well illustrated by our results on *D. melanogaster*, showing that three paralogous genes on the X chromosome contribute strongly to the average positive LD measured among deleterious mutations, due to the presence in some individuals of a diverged sequence that is absent from the reference genome and mapped on one of these paralogs. As discussed above, this problem cannot be solved by eliminating variants categorized as deleterious and present above a threshold frequency, as this would automatically generate a bias towards positive LD. Ideally, pseudo-heterozygous SNPs should be removed from the analysis, but their detection is extremely challenging in an outcrossing diploid species. Our results on *C. grandiflora* show that removing about 150 small genomic regions in which at least 5 alleles categorized as deleterious are only present in the heterozygous state (in the same individuals for all alleles) and where coverage is significantly higher in heterozygous individuals already reduces substantially the average LD among deleterious mutations. However, our criterion is probably conservative, and the exact contribution of pseudo-heterozygous sites to average LD is difficult to assess. Besides, inferring the structural variants at the origin of these mapping errors is challenging with short-reads. Indeed, using Smoove and Delly to validate blocks of heterozygosity we found that only a fraction of them show a clear overlap with duplications but that most SV calls and coverage pattern were inconsistent and noisy (results not shown). The generalization of long-read sequencing datasets will certainly open new possibilities for estimating LD, being much less prone to this type of artifact (De Coster et al., 2021; Igolkina et al., 2025).

In the absence of any conditioning on allele frequency, and if mapping errors and other types of possible artifacts could be ruled out with a high level of confidence, the persistence of average positive LD among deleterious mutations would imply that a selective mechanism generating LD must be involved, if the average LD among neutral mutations stays close to zero (as in the datasets analyzed here). Different sources of positive LD are possible. A first, already considered by other authors (Ragsdale, 2022; Sandler et al., 2021; Sohail et al., 2017; Stolyarova et al., 2022) is positive epistasis among deleterious alleles. This may correspond to possible compensatory effects between mutations within the same gene (Davis et al., 2009), or to the fact that, once a gene has become non-functional due to a first mutation, additional mutations within the same gene become neutral and can accumulate. Some evidence for possible compensatory effects of mutations is given in Ragsdale (2022) and Stolyarova et al. (2022). In particular, without conditioning on allele frequencies, Ragsdale (2022) found positive LD at very short distances (<300 bp) inside genes, for both missense and synonymous mutations in several human populations. No difference in LD was observed between synonymous and missense mutations when averaging within genes, but LD was higher between missense mutations inside conserved gene domains compared to synonymous mutations (while the opposite pattern was found outside conserved gene domains), which was interpreted as evidence for positive epistasis among deleterious mutations affecting the same conserved domain. Furthermore, Stolyarova et al. (2022) showed that, controlling for the physical distance in base pairs among mutations, higher positive LD is observed among mutations affecting amino acids that are physically close to each other in the protein 3D configuration than among mutations affecting amino acids that are farther apart, in two populations of the fungus *Schizophyllum commune*. Johri and Charlesworth (2025) recently showed that another possible source of positive LD among deleterious alleles corresponds to the non-complementation of mutations within the same gene in diploids. Under this scenario, the fitness of doubly heterozygous individuals in which each gene copy carries a different mutation is lower than the fitness of doubly heterozygous individuals in which both mutations are present in the same copy, due to the fact that the second type of individuals maintains a fully functional gene copy. This may be seen as a particular case of epistasis between mutations in trans, whose effect (generating positive LD) is generally weaker than the effect of epistasis between mutations in cis (Johri and Charlesworth, 2025).

Positive LD among deleterious alleles may also result from the mixing of haplotypes from subpopulations in which the frequencies of these alleles differ. This can occur in spatially structured populations when the strength of selection against deleterious alleles is stronger in some subpopulations than in others (Roze, 2012), leading to a lower frequency of deleterious alleles on haplotypes from subpopulations under stronger selection. These differences in selection may stem from environmental differences (for example, if selection against deleterious alleles is stronger in harsher environments, Agrawal and Whitlock, 2010), or from stronger effects of drift in some subpopulations (that may result from range expansion dynamics, e.g., Peischl et al., 2013) leading to an increased frequency of deleterious alleles. This type of process is unlikely to be a cause of positive LD among deleterious alleles in the *C. grandiflora* and *D. melanogaster* datasets analyzed here, since individuals were sampled from single populations with low structure (Lack et al., 2015; Sandler et al., 2021; St. Onge et al., 2011), but may be a possible source of positive LD in other datasets. Similarly, positive LD among deleterious alleles may be caused by a difference in the strength of selection against those alleles between the sexes (Lenormand, 2003), which may result from stronger sexual selection among males (Whitlock and Agrawal, 2009). Finally, positive LD may be generated by differences among diploid individuals in their degree of inbreeding. Indeed, more inbred lineages tend to be relatively more homozygous than less inbred lineages, and therefore purge deleterious alleles more efficiently, leading to a lower frequency of deleterious alleles. In particular, this effect is a deterministic source of positive LD in partially self-fertilizing populations, in the absence of epistasis among selected loci (Roze and Lenormand, 2005; Stetsenko and Roze, 2022). The same effect also occurs in finite or spatially structured diploid populations due to random variations on the degree of inbreeding of individuals. In this case, the sign of LD among deleterious alleles depends on the relative effects of the variance in inbreeding generating positive LD, and of the Hill-Robertson effect generating negative LD (Fouqueau and Roze, 2025; Roze, 2021). In a single finite population and when deleterious alleles stay near mutation-selection balance, one predicts that LD among deleterious alleles should be positive when their dominance coefficient *h* is less than 0.25, and negative when *h >* 0.25 (Roze, 2021).

The continuous advancement of DNA sequencing technologies is opening exciting new avenues for disentangling the causes of associations between alleles at selected loci within natural populations. This progress may, in turn, shed light on the evolutionary consequences of deleterious mutations on genetic architecture and reproductive systems. However, it also presents significant methodological challenges, as estimates can be strongly influenced by technical artifacts. Enhancing the robustness and accuracy of inference methods therefore represents a promising direction for future research.

## Methods

### Coalescent simulations

The effect of computing LD and other two-locus moments on subsets of neutral variants present in a given frequency class in a sample of *n* sequences was checked using coalescent simulations. We simulated the ancestral recombination graph (ARG) of a sample of sequences (each with an infinite number of sites at which mutations can occur) using Hudson’s algorithm (p. 139-144 in Hein et al., 2004) with parameters *ρ* = 4*Nr* and *θ* = 4*Nu*, where *N* is population size, *r* the recombination rate (probability that a crossover occurs within the sequence) and *u* the mutation rate within the sequence (*ρ* = 40 and *θ* = 100 were used for all results presented here, while sample size was set to *n* = 400 since the dataset from *C. grandiflora* has approximately 200 diploid individuals). Pairs of segregating sites were classified into 100 classes of genetic distance, and the linkage disequilibrium *D_ab_* among derived alleles (and other two-locus moments) was averaged over 20 batches of 500 coalescent trees (10^4^ trees in total). 95% confidence intervals of the ratio ⟨*D_ab_*⟩ */* ⟨*p_a_q_a_p_b_q_b_*⟩ were obtained using the bootstrap method for each distance class, by resampling 1000 times over batch averages. In order to assess the effect of ancestral state misspecification, we also included a parameter *µ* corresponding to the probability that a variant is misspecified.

### Population genomic data

We retrieved the whole-genome sequencing data from 182 individuals of the obligate outcrossing plant *Capsella grandiflora* from Northern Greece (Josephs et al., 2015) that was used by Sandler et al. (2021). This dataset is suitable for testing predictions on LD patterns from classical population genetics model, as it is composed of a large number of individuals sampled from a single population with low genetic structure of an obligately outcrossing species, limiting the effects of population structure and inbreeding. The list of individuals is available in Table S1. Raw reads were trimmed using Trimmomatic v0.39 (Bolger et al., 2014) and mapped on a new (not yet published) genome of *Capsella rubella*, Cr145 (Tianlin Duan, personal communication) using bwa v0.7.17 (Li and Durbin, 2010). The Cr145 genome has a total length of 158,571,049 bp divided into 129 contigs with an N50 of 17,600,677 bp. The mean coverage per individual was 39.99X. Duplicate reads where marked using the *MarkDuplicates* option of Picard Tools v2.23.5 (broadinstitute.github.io/picard/) and removed from subsequent analysis. We obtained a VCF file containing 20,745,523 called sites between all individuals. Variants were polarized using an alignment of the whole genome assembly of the related Brassicaceae *Neslia paniculata* (Slotte et al., 2013) onto the *C. rubella* Cr145 genome (Tianlin Duan, personal communication). The *Drosophila melanogaster* data consisted in the VCF file obtained by Sandler et al. (2021), corresponding to SNP data from 190 haploid embryos from the Zambian population DPGP3 (Lack et al., 2015), mapped onto the reference genome 5.57 of *D. melanogaster* (file ZI_allchrom_clean_bi_samples.ann.vcf downloaded from https://zenodo.org/records/4895505). Variants were polarized using an alignment of the whole genome assembly of *Drosophila yakuba* (release 1.3) onto the the reference genome 5.57 of *D. melanogaster* obtained using LASTZ and Kent utilities (see pipeline in github.com/RomanStet/LD_del_mut).

### SNP calling

Single nucleotide polymorphisms (SNPs) of *C. grandiflora* were called using GATK and following the GATK Best Practices (Van der Auwera et al., 2013) and adjustments were made to the filtering criteria in the *HaplotyeCaller* option based on the distribution of quality scores with filters set to: *MQ<50, SOR>3, QD<2, FS>60, MQRankSum<-5, ReadPosRankSum<-8, ReadPosRank-Sum>8*. In addition, we filtered out sites with extreme mean coverage (>71.65X) and genotypes with low mean coverage (<10X; Figure S23). This was done to filter out the lowest confidence genotypes and regions with a high number of reads due to repetitive sequences. Monomorphic sites, sites with more than 2 alleles and sites with a high proportion of missing data (>30%) were also filtered out to obtain 8,494,402 biallelic SNPs. Based on a PCA with the filtered independent SNPs (using the function *indep-pairwise* of Plink with parameters: window size = 50 kb, step size = 100 variants and *r*^2^ threshold = 0.1; Purcell et al., 2007), we excluded from LD computation 11 genetically divergent individuals (Figure S24, Table S1). Most of these divergent individuals also present outlier proportions of missing data (Figure S25).

### SIFT annotation

The SIFT4G algorithm (Sorting Intolerant From Tolerant For Genomes, Vaser et al., 2016) was used to categorize mutations based on their potential fitness effect. This program uses multiple protein alignments across a database of proteins from a large diversity of taxa, to categorize each possible nucleotide at each coding site as deleterious or tolerated. If the site is conserved across the alignment and the query nucleotide is present in a low number of sequences it is inferred as deleterious whereas if the site is not conserved across the alignment or the query nucleotide is present in a higher proportion of sequences it is inferred as tolerated. More precisely, a score from 0 to 1 is attributed to each possible nucleotide at each site, and SIFT4G categorizes nucleotides with a SIFT score ≤ 0.05 as deleterious and *>* 0.05 as tolerated. We built a SIFT4G library for the Cr145 *C. rubella* reference genome and the *D. melanogaster* 5.57 reference genome using the UniRef90 protein database (uniprot.org/uniref/). In order to distinguish between potentially mildly deleterious mutations from more strongly deleterious mutations, we split the category of nucleotides with a SIFT score lower than 1 into “mildly deleterious mutations” (0.05 < SIFT score < 1) and “deleterious mutations” (SIFT score ≤ 0.05).

### Detecting potential structural variants in *C. grandiflora*

Errors may occur during the mapping of short-read sequencing data, which may cause patterns of pseudo-heterozygosity and give rise to spurious LD. This was shown in particular using data from the 1001 Arabidopsis Genome Project, in which a high proportion of heteroygous SNPs were identified, while *A. thaliana* is highly selfing (Jaegle et al., 2023). As shown by Jaegle et al., 2023, mapping errors may result from polymorphic duplications. Indeed, if a duplication is present in a subset of individuals only and absent from the reference genome, reads from the duplicated sequence will be mapped onto the original sequence, and mutations present in the duplicated sequence will thus give rise to spurious heterozygosity at the corresponding sites in the original sequence (Figure S17). In principle, the *D. melanogaster* dataset should be much less affected by this type of error, since sequences were obtained from haploid embryos and pseudo-heterozygous sites have been discarded (Lack et al., 2015).

We wrote an R script to detect potential pseudo-heterozygosity in the *C. grandiflora* dataset, that may be caused by polymorphic duplications (or other possible sources of mapping errors). Indeed, such duplications may be detected from neighboring sites presenting similar patterns of heterozygosity (*i.e.*, the same individuals are heterozygous). Mutations that are fixed in the duplicated sequence but absent from the original sequence will thus always appear in the heterozygous state (Figure S17, case 1), while mutations that are fixed in the original sequence but absent from the duplicated sequence will appear in the heterozygous state in the same individuals as case 1 mutations, while they will appear in the homozygous state in all other individuals (Figure S17, case 2). Our R script thus uses data from the SNP calls of *C. grandiflora* to detect blocks of genome carrying sites that present similar patterns of heterozygosity, and at which only two genotypes are present in the sample: heterozygous and homozygous for the ancestral allele (case 1), or heterozygous and homozygous for the derived allele (case 2). In order to take into account mutations that may not be fixed within the duplicated or original sequence (and genotyping errors), two sites falling either into case 1 or case 2 were considered as potentially indicative of a polymorphic duplication when they shared at least 70% of their heterozygous individuals. Sites are examined sequentially along the chromosome until no case 1 or case 2 pair of sites sharing a common pattern of heterozygosity is found over a distance of 2 kb. Among the large number of genome blocks detected, only a minority may correspond to polymorphic duplications (in particular, the majority of detected blocks only contain 2 or 3 sites sharing a common pattern of heterozygosity). Furthermore, our algorithm should detect haplotype blocks present at low frequency (since these blocks will tend to carry mutations that are mostly present in the heterozygous state) that do not necessarily correspond to potential duplications. We thus only retained blocks in which at least five deleterious mutations shared a common pattern of heterozygosity, and for which the coverage (over the detected block) in heterozygous individuals that potentially carry the duplication (scaled by the mean coverage of the individual) is significantly and at least 1.3 times higher than the scaled coverage in the other individuals.

This list of blocks of heterozygosity in *C. grandiflora* was compared to outputs of two standard structural variant (SV) detection programs for short read sequencing, Smoove and Delly (Rausch et al., 2012). Smoove is based on the LUMPY algorithm (Layer et al., 2014), and like Delly, uses information on paired-ends, split-reads and read-depth to detect SV. We ran both programs on mapped reads (see *SNP calling* above) for each individual of *C. grandiflora* and retained only SV calls not tagged as ‘low quality’ by Smooove and Delly and shorter than 5 Mbp to filter out spurious SV calls. When an SV call from Smoove or Delly overlapped with a block of the list obtained using our R script described above, we computed the ratio of their lengths (length ratio) and the ratio of the length of the overlapping portion over the length of the longer element (overlap ratio). The overlap ratio equals 1 if the SV call and the block completely overlap and approaches 0 if the length of the overlapping portion becomes negligible compared to the longest element. We retained cases in which the overlap ratio was larger than 0.05. We then used Samplot (Belyeu et al., 2021) to visualize SV calls and read depth in the region of the block for all heterozygous individuals and 5 randomly drawn homozygous individuals. We considered that the heterozygosity block was validated (either by Smoove or Delly) if SVs were only called in heterozygous individuals and if the read depth was changed at the location of the SV.

### Computing LD

Since SNP data from *C. grandiflora* are unphased, the two double heterozygous genotypes *Ab/aB* and *ab/AB* cannot be distinguished. Therefore, we measured LD between alleles *a* and *b* using the composite LD measure Δ*_ab_*, which is the sum of the association between *a* and *b* on the same chromosome (*D_ab_*) and the association between *a* and *b* on different chromosomes (*D_a,b_* in the notation of Kirkpatrick et al., 2002). This composite LD can be computed as:

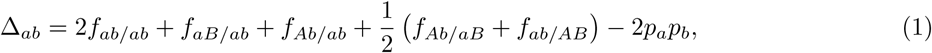

where *f_X_* is the frequency of genotype *X* in the sample, and *p_a_*, *p_b_* the frequencies of *a* and *b* (e.g., p. 126 in Weir, 1996). In the case of tightly linked loci, *D_a,b_* ≈ *F D_ab_*, where *F* is the inbreeding coefficient (Nordborg, 1997; Roze, 2016), so that Δ*_ab_* ≈ *D_ab_* (1 + *F*). Therefore, Δ*_ab_* should be equal to *D_ab_* under random mating, and is increased by inbreeding. Since the *D. melanogaster* dataset was obtained from haploid embryos, LD between variants present on the same chromosome (*D_ab_*) was computed for this datasets.

LD between neutral variants was computed from sites at which segregating nucleotides have both a SIFT score of 1. In this case, *a* and *b* correspond to the derived alleles based on the sequence of the outgroup (*N. paniculata* for *C. grandiflora* and *D. yakuba* for *D. melanogaster*). In *C. grandiflora* and *D. melanogaster*, LD between deleterious alleles was computed from sites at which one segregating nucleotide has a SIFT score of 1, and the other a SIFT score ≤ 0.05: in this case, *a* and *b* correspond to the alleles with SIFT score ≤ 0.05 (thus putatively deleterious). Finally, in both species, LD between mildly deleterious alleles was computed from sites at which one segregating nucleotide has a SIFT score of 1, and the other a SIFT score between 0.05 and 1 (and where *a* and *b* are the alleles with SIFT score between 0.05 and 1). We checked that restricting the analyses of LD between deleterious or mildly deleterious alleles on derived alleles did not affect the results: indeed, most mutations with a SIFT score < 1 are derived (Figure S13). The mean value of Δ*_ab_* and *D_ab_* was computed for every pair of sites at different physical distances up to 1.5 kb (each distance class being 25 bp wide), discarding pairs of sites for which the information on genotype was not available (at both sites) in more than half of the total number of individuals. As for the LD estimator *σ*^2^ of Ohta and Kimura (1969), the average LD for a given distance class ⟨Δ*_ab_*⟩ or ⟨*D_ab_*⟩ was scaled by the average product of allele diversities at both loci (for the same distance class) ⟨*p_a_q_a_p_b_q_b_*⟩, where ⟨*X*⟩ denotes the average of quantity *X* over pairs of sites in the same distance class. 95% confidence intervals were computed using the bootstrap method, by resampling each set of pairs of sites 1,000 times.

## Acknowledgments

We thank Tianlin Duan for sharing the reference genome of *C. rubella*. We thank the bioinformatics and computing service of Roscoff’s Biological Station (Abims platform) for computing time. We also thank Aneil Agrawal for helpful comments, and Guillaume Achaz for very useful discussions on the coalescent tree interpretation of LD between neutral mutations.

## Data availability

Scripts used in the bioinformatic analyses are available on github.com/RomanStet/LD_del_mut.

## Supplementary figures

**Figure S1:**
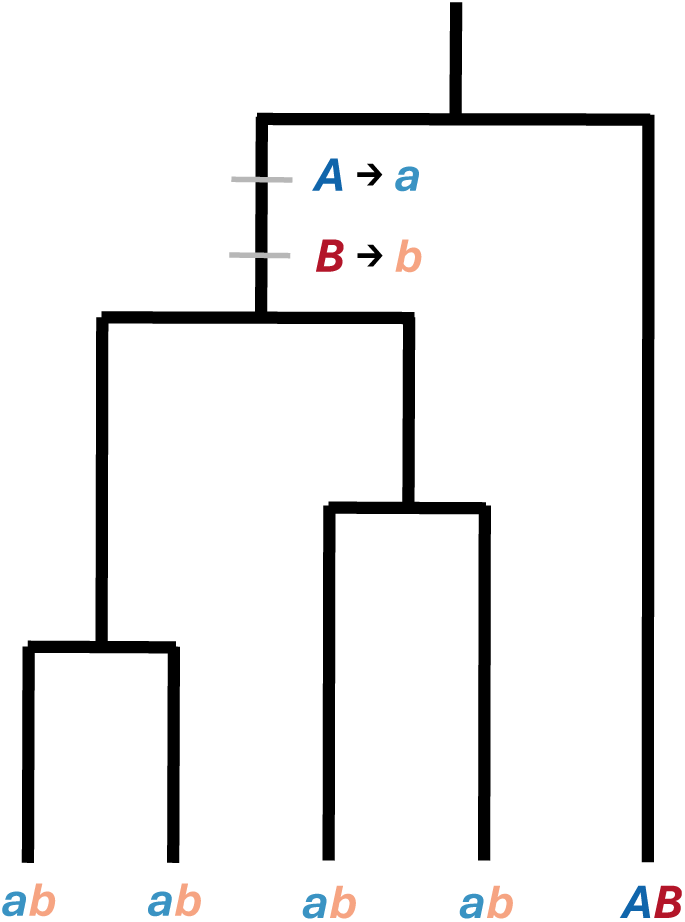
Example of a coalescence tree where derived alleles *a* and *b* are at the same frequency ((*n* — 1)*/n*) in a sample of size *n* = 5. If no recombination is assumed between the two loci, the two mutation events (from *A* to *a* and from *B* to *b*) must have arisen in the same part of the tree (between the coalescence event of all *a* and all *b* alleles and the last common ancestor of the entire sample). Note that, in this case, alleles *a* and *b* are in positive LD (*D_ab_ =* 0.16).

**Figure S2:**
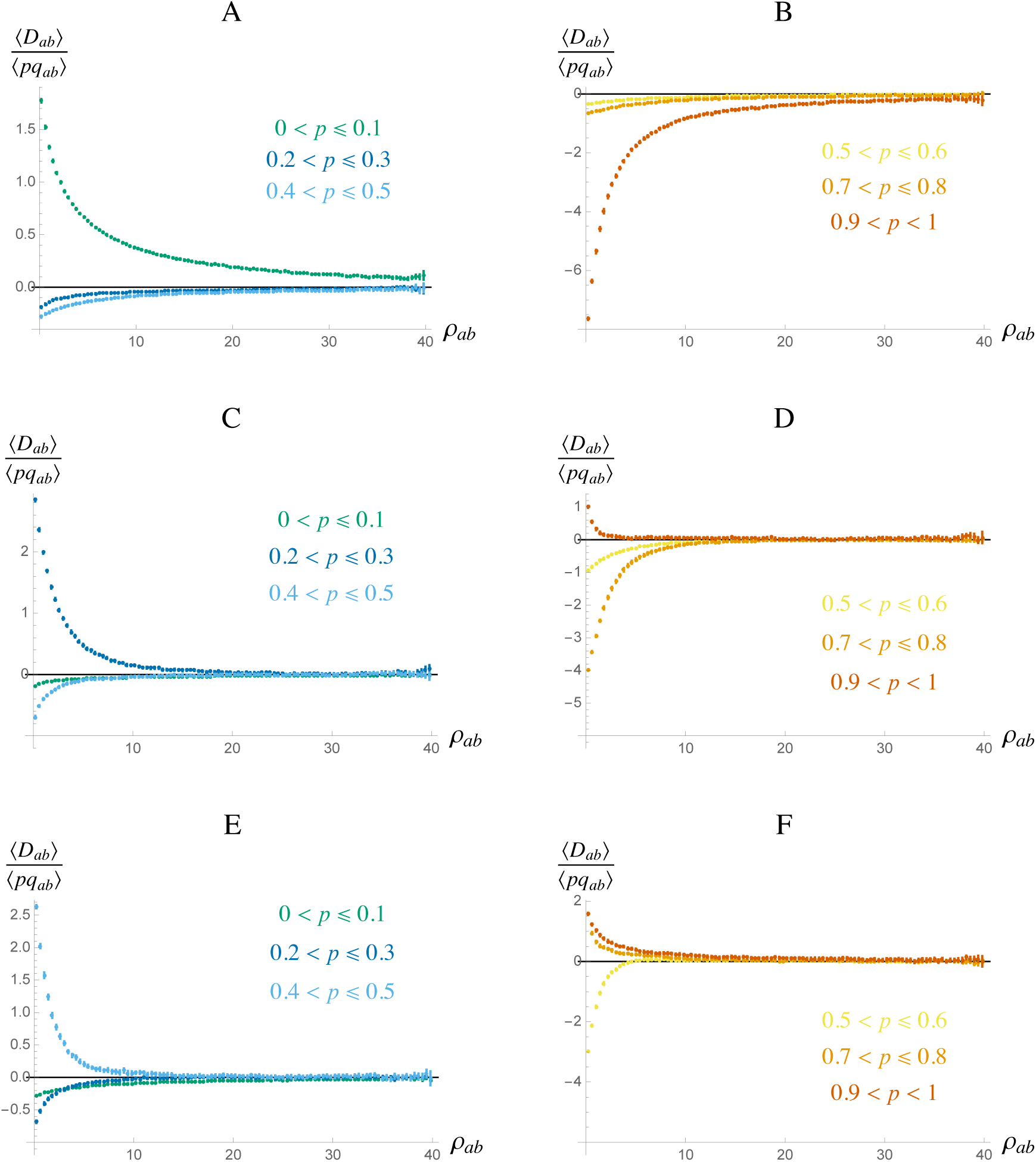
Average LD ⟨*D_ab_*⟩ between derived neutral mutations (scaled by the average product of genetic diversities ⟨*pq_ab_*⟩ = ⟨*p_a_q_a_p_b_q_b_*⟩) in the coalescent simulations, as a function of the recombination rate *ρ_ab_* = 4*N_e_r_ab_* between mutations with different frequencies. LD is averaged over pairs of mutations, the first mutation belonging to a given frequency class (0 *< p* ≤ 0.1 for A and B; 0.2 *< p* ≤ 0.3 for C and D; 0.4 *< p* ≤ 0.5 for E and F) and the second mutation to the frequency class shown by the colors. Errors bars correspond to 95% confidence intervals (see Methods section). Parameters values: *θ* = 4*N_e_u* = 100 (mutation rate); *n* = 400 (sample size).

**Figure S3:**
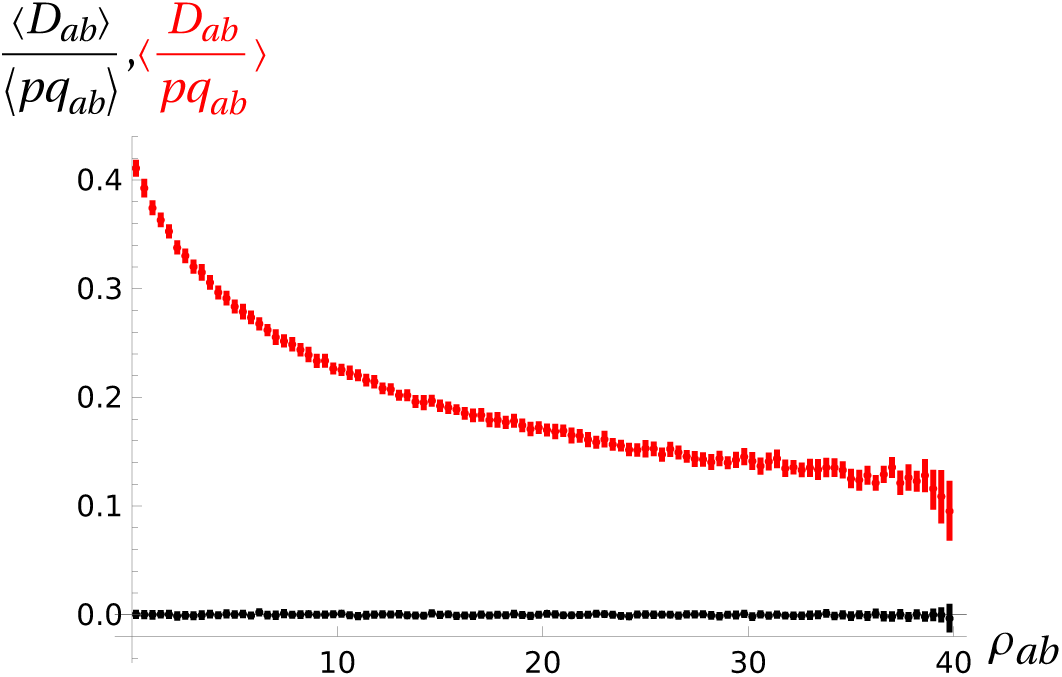
Average LD ⟨*D_ab_*⟩ between derived neutral mutations (scaled by the average product of genetic diversities (*pq_ab_*) = (*p_a_q_a_p_b_q_b_*); black) or average of *D_ab_/pq_ab_* (red) over all segregating sites (no conditioning on frequency) in the coalescent simulations, as a function of the recombination rate *ρ_ab_* = 4*N_e_r_ab_*. Errors bars correspond to 95% confidence intervals (see Methods section). Parameters values: *θ =* 4*N_e_u* = 100 (mutation rate); *n* = 400 (sample size).

**Figure S4:**
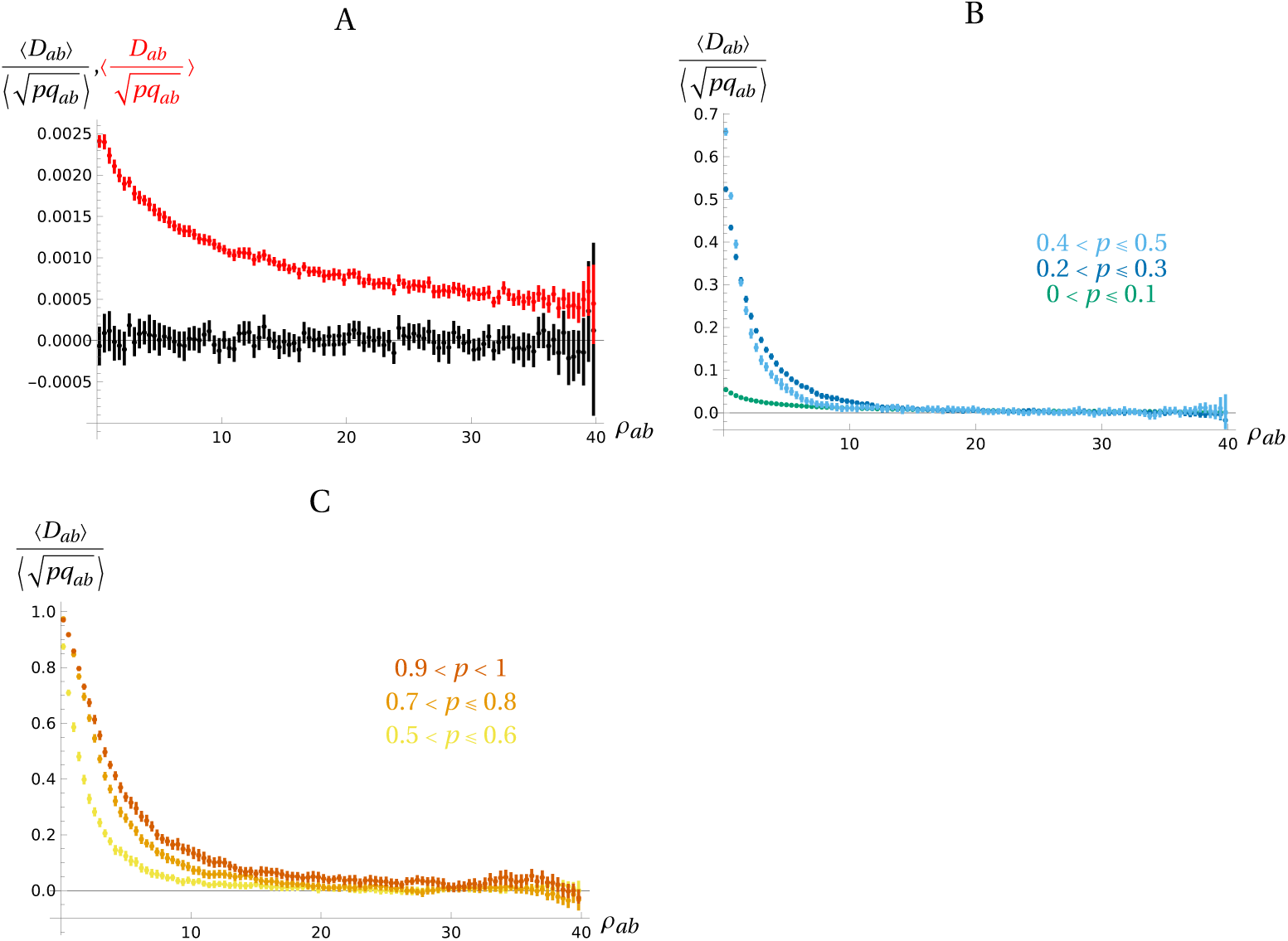
Average LD ⟨*D_ab_*⟩ between derived neutral mutations (scaled by the average square root of the product of genetic diversities 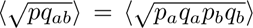) in the coalescent simulations, as a function of the recombination rate *ρ_ab_* = 4*N_e_r_ab_*• A: Average of *D_ab_* (scaled by the average of 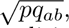 black) and average of 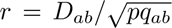 (red) over all segregating sites for each distance class (no conditioning on frequency). B and C: Average of *D_ab_* (scaled by the average of 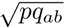) over pairs of loci at which the derived allele segregates in a given frequency range. Errors bars correspond to 95% confidence intervals (see Methods section). Parameters values: *θ* = 4*N_e_u* = 100 (mutation rate); *n* = 400 (sample size).

**Figure S5:**
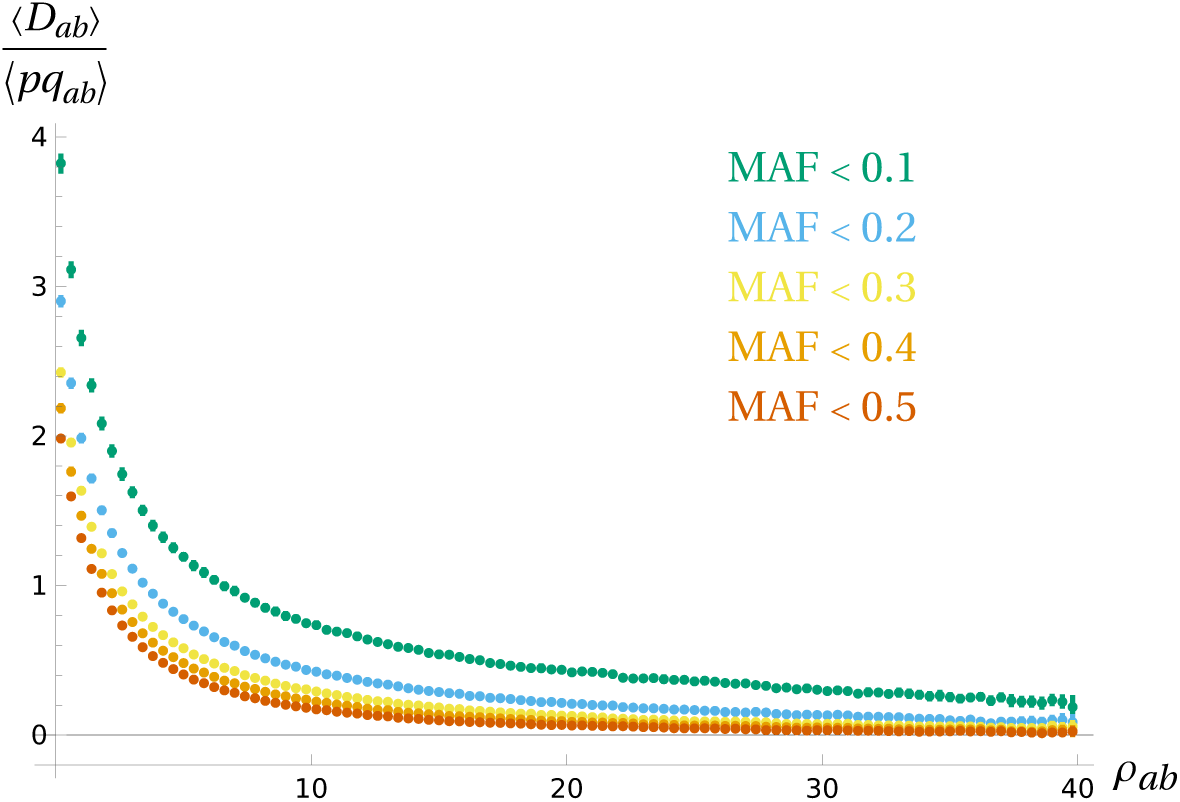
Average LD ⟨*D_ab_*⟩ between neutral mutations (scaled by the average product of genetic diversities (*pq_ab_*) = (*p_a_q_a_p_b_q_b_*)) in the coalescent simulations, as a function of the recombination rate *ρ_ab_ =* 4*N_e_r_ab_*. LD is polarized based on the minor allele frequency (MAF) and averaged over pairs of sites at similar distances, for different MAF thresholds. Parameter values are as in Figure 1.

**Figure S6:**
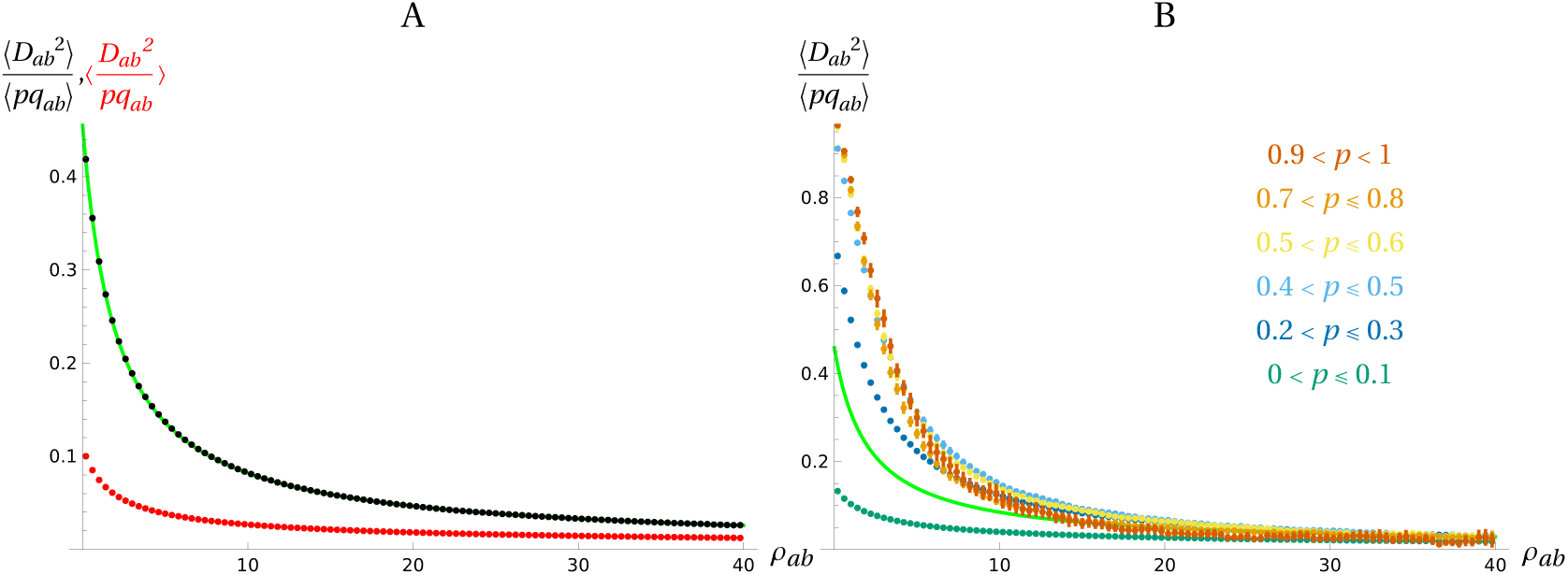
Squared LD ⟨*D_ab_*^2^⟩ between derived neutral mutations (scaled by the average product of genetic diversities (*pq_ab_*) = (*p_a_q_a_p_b_q_b_*)) in the coalescent simulations, as a function of the recombination rate *ρ_ab_ =* 4*N_e_r_ab_*. A: Average of *D_ab_*^2^ (scaled by the average of *pq_ab_*, black) and average of *D_ab_*^2^*/pq_ab_* (red) over all segregating sites for each distance class (no conditioning on frequency). B: Average of *D_ab_*^2^ (scaled by the average of *pq_ab_*) over pairs of loci at which the derived allele segregates in a given frequency range. The green solid curves correspond to the analytical prediction (10 + *ρ_ab_*) */* [(2 + *ρ_ab_*) (11 + *ρ_ab_*)] from Ohta and Kimura (1969) when averaging over all segregating sites. Parameters values are as in Figure 1.

**Figure S7:**
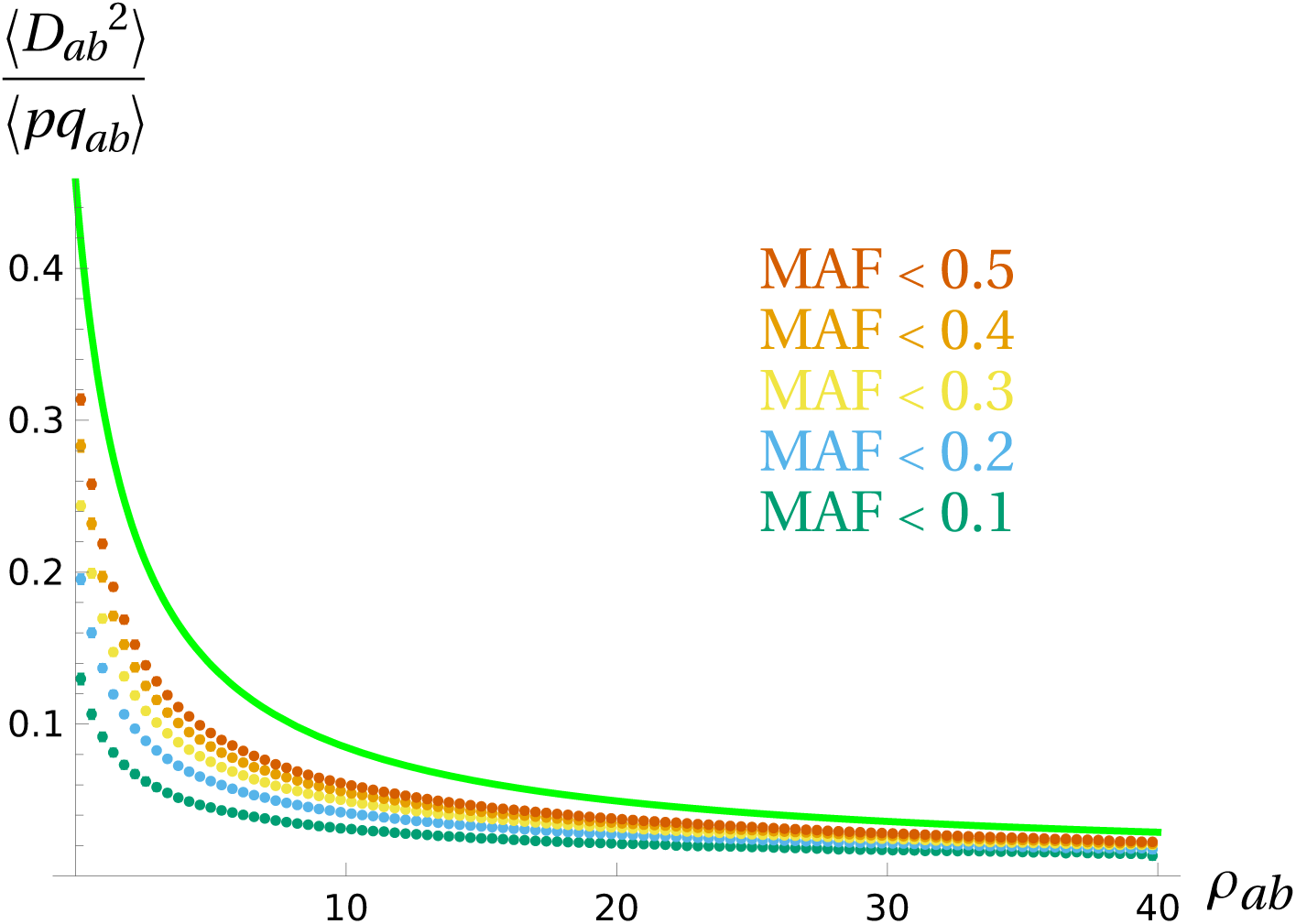
Squared LD ⟨*D_ab_* ^2^⟩ between neutral mutations (scaled by the average product of genetic diversities (*pq_ab_*) = (*p_a_q_a_p_b_q_b_*)) in the coalescent simulations, as a function of the recombination rate *ρ_ab_* = 4*N_e_r_ab_*. LD is polarized based on the minor allele frequency (MAF) and averaged over pairs of sites with different MAF thresholds. The green solid curve correspond to the analytical prediction (10 + *ρ_ab_*) */* [(2 + *ρ_ab_*) (11 + *ρ_ab_*)] from Ohta and Kimura (1969) when averaging over all segregating sites. Parameter values are as in Figure 1.

**Figure S8:**
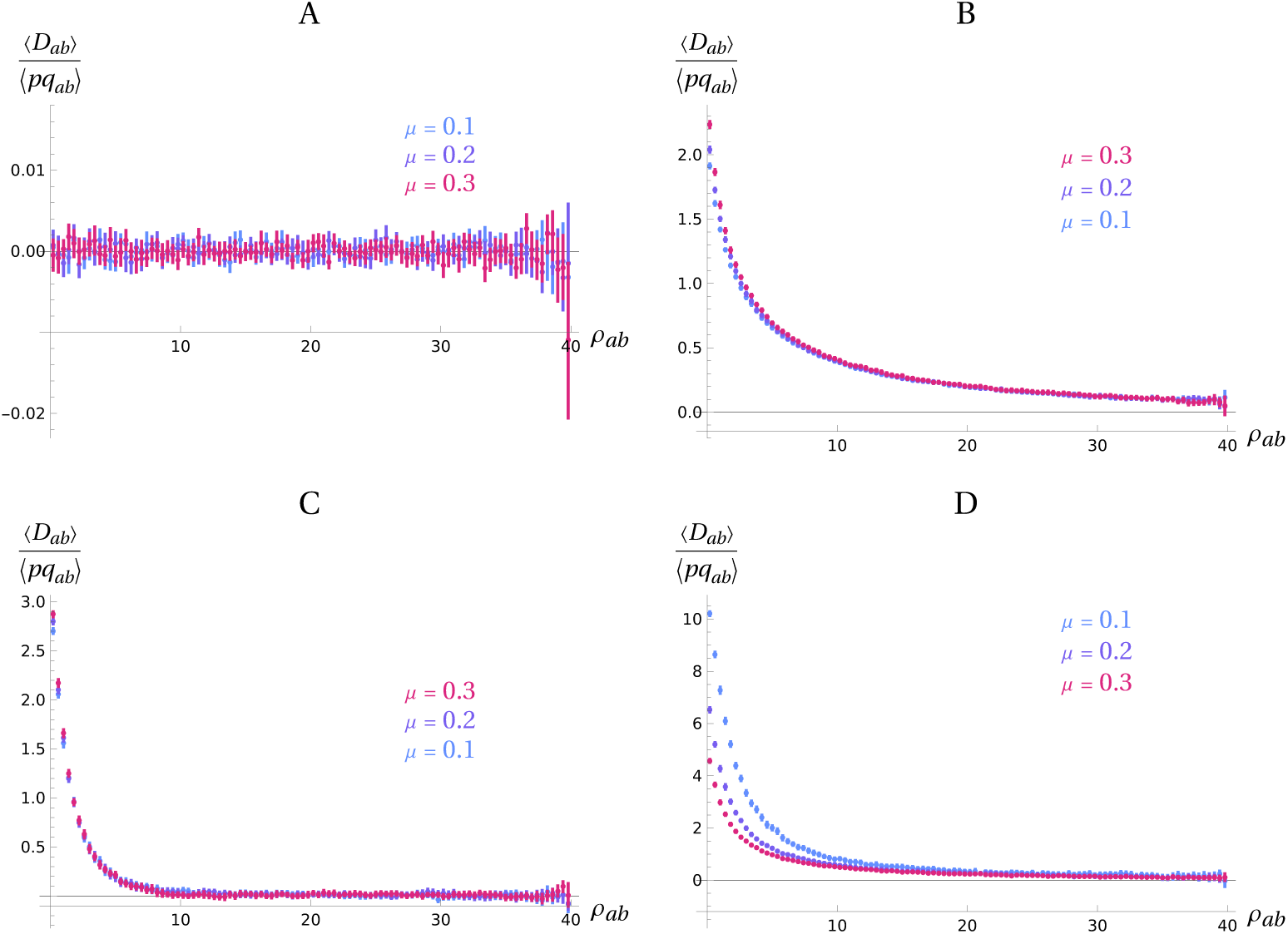
Average LD ⟨*D_ab_*⟩ between derived neutral mutations (scaled by the average product of genetic diversities ⟨*pq_ab_*⟩ = ⟨*p_a_q_a_p_b_q_b_*⟩) in the coalescent simulations, as a function of the recombination rate *ρ_ab_* = 4*N_e_r_ab_*, and for different values of the rate of misspecification *µ* of the ancestral/derived state of alleles. A: Averages over all segregating sites for each distance class (no conditioning on frequency); B: only the lowest frequency class (*p* ≤ 0.1, green points in Figure 1A); C: only an intermediate frequency class (0.4 *< p* ≤ 0.5, light blue points in Figure 1A); D: only the highest frequency class (*p >* 0.9, dark orange points in Figure 1B). Parameter values are as in Figure 1.

**Figure S9:**
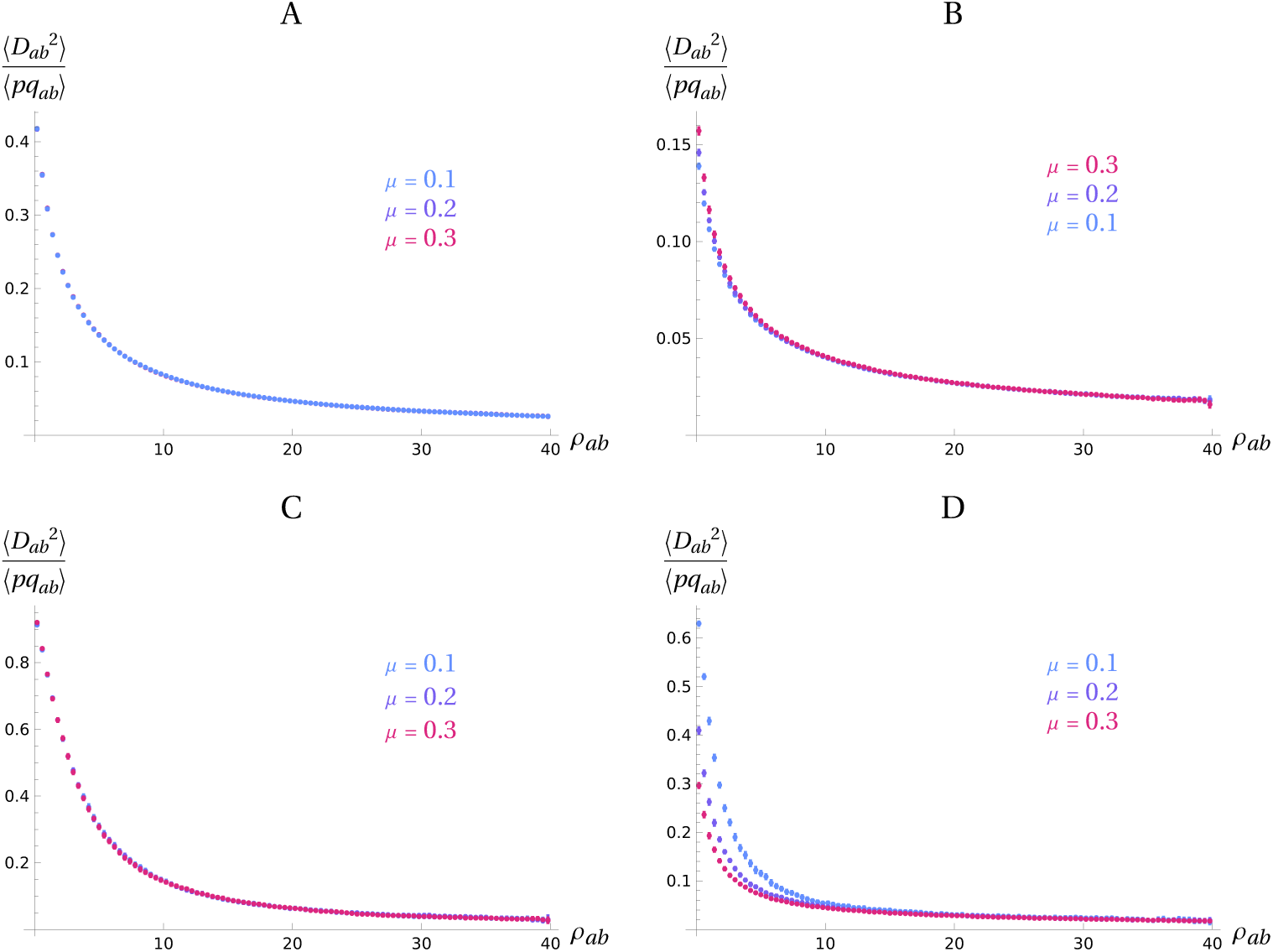
Squared LD ⟨*D_ab_*^2^⟩ between derived neutral mutations (scaled by the average product of genetic diversities ⟨*pq_ab_*⟩ = ⟨*p_a_q_a_p_b_q_b_*⟩) in the coalescent simulations, as a function of the recombination rate *ρ_ab_* = 4*N_e_r_ab_*, and for different values of the rate of misspecification *µ* of the ancestral/derived state of alleles. A: Averages over all segregating sites for each distance class (no conditioning on frequency); B: only the lowest frequency class (*p* ≤ 0.1, green points in Figure S6B); C: only an intermediate frequency class (0.4 *< p* ≤ 0.5, light blue points in Figure S6B); D: only the highest frequency class (*p >* 0.9, dark orange points in Figure S6B). Parameter values are as in Figure 1.

**Figure S10:**
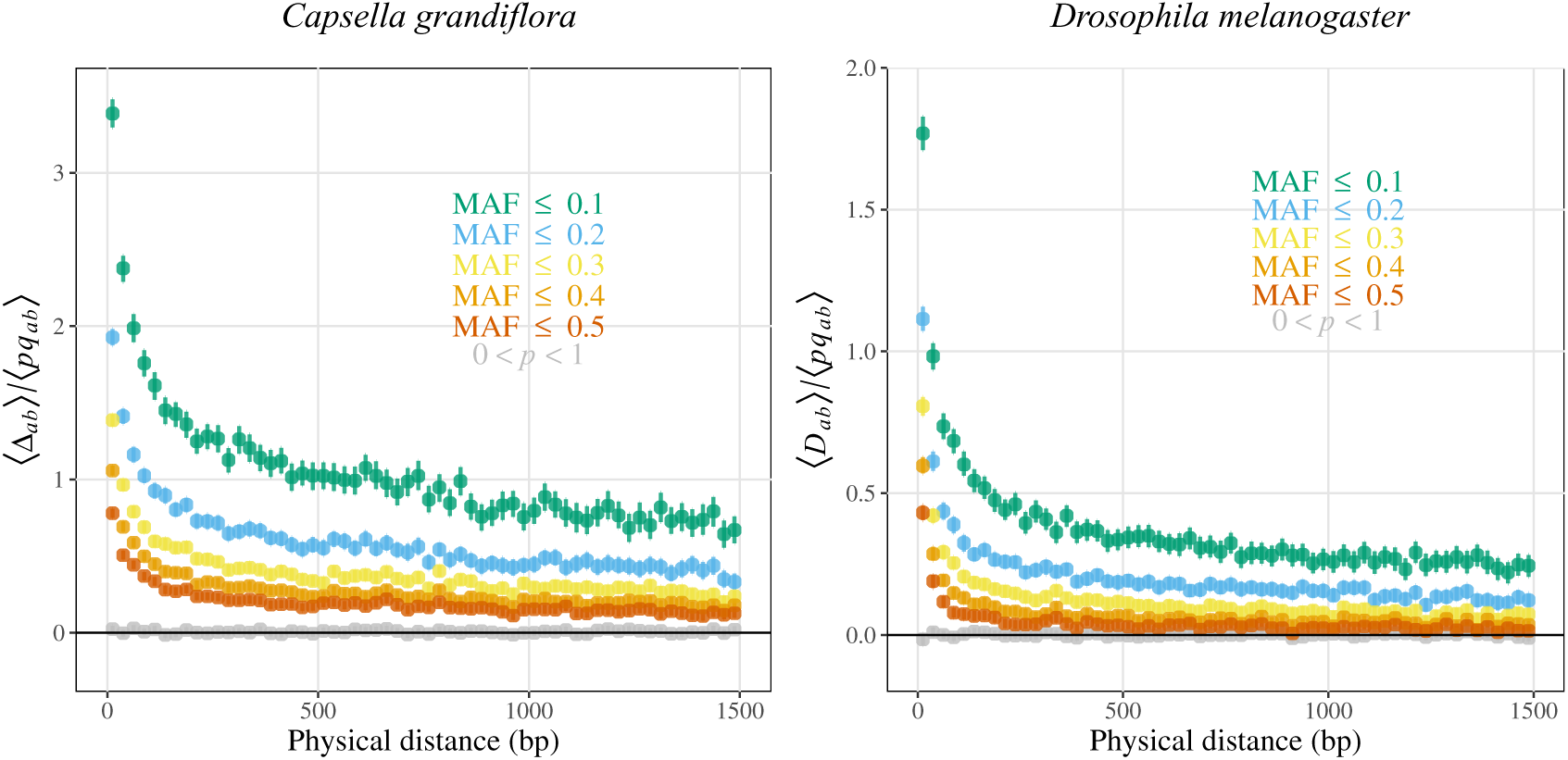
Average composite linkage disequilibrium Δ*_ab_* or linkage disequilibrium *D_ab_* (scaled by the average product of allele frequencies) between neutral derived variants according to physical distance between variants, in *C. grandiflora* and *D. melanogaster*. In color: LD is polarized based on the minor allele frequency (MAF) and averaged over pairs of sites at similar distances, for different MAF thresholds; In grey: LD is computed between neutral derived variants irrespective of their frequency and averaged over pairs of sites at similar distances.

**Figure S11:**
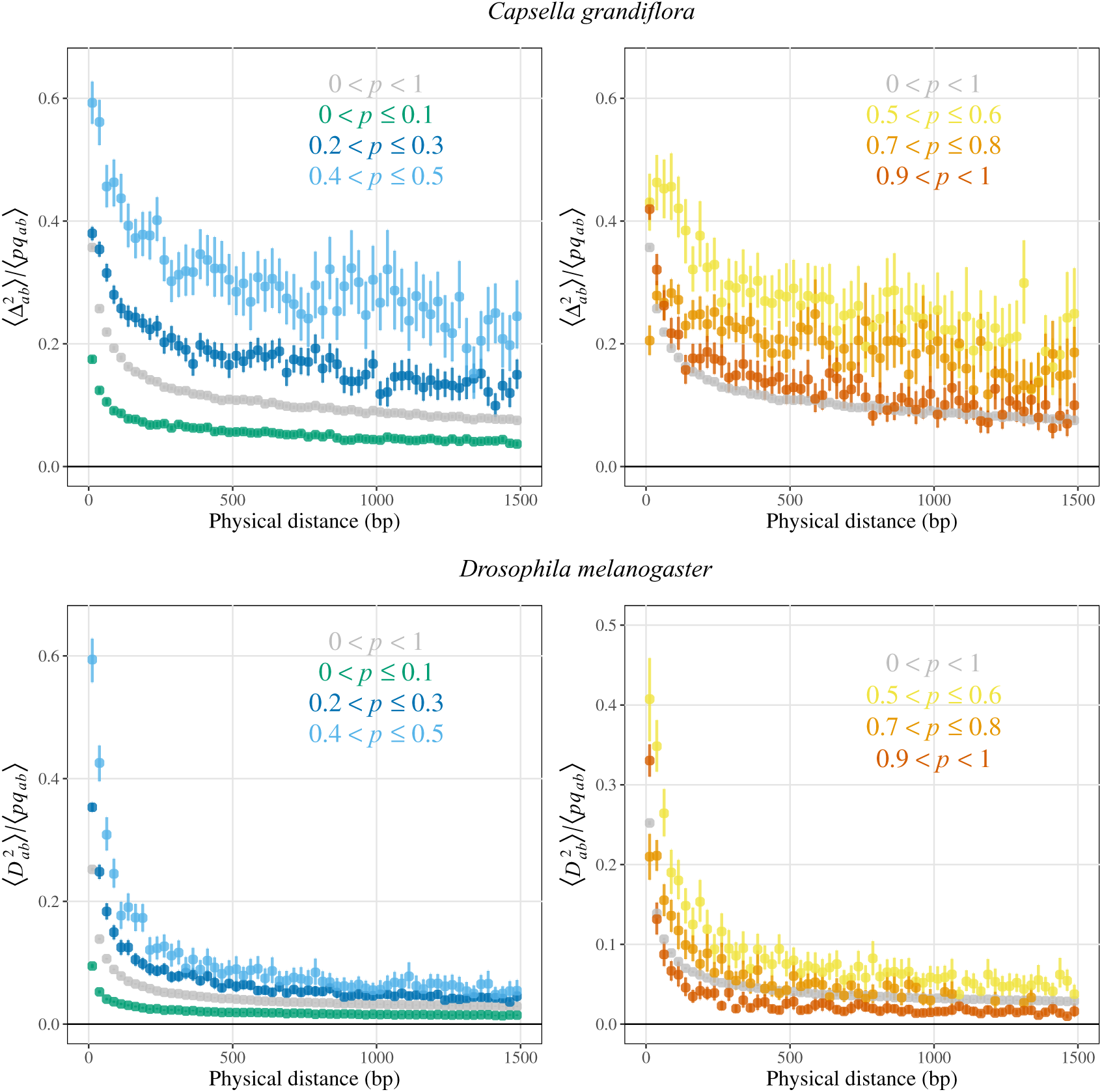
Average squared composite LD Δ_ab_^2^ or squared LD *D_ab_*^2^ between neutral derived variants (scaled by the average product of allele frequencies) according to physical distance between variants, in C. grandiflora and D. melanogaster. In color: LD is computed over pairs of loci at which the derived allele segregates in a given frequency range (same as in Figure 1C and 1D); In grey: LD is computed between neutral derived variants irrespective of their frequency.

**Figure S12:**
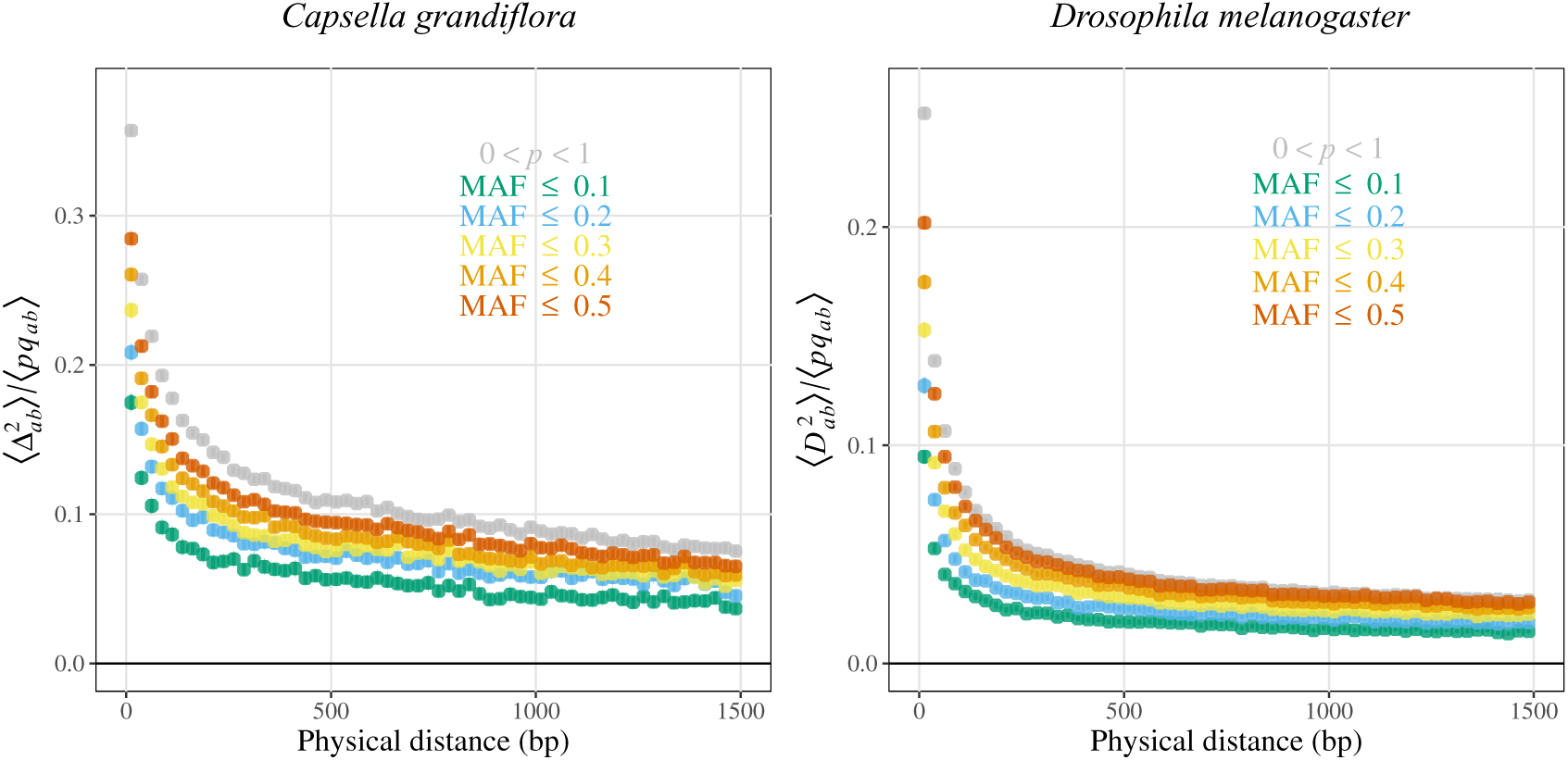
Average squared composite LD Δ*_ab_*^2^ or squared LD *D_ab_*^2^ between neutral mutations (scaled by the average product of allele frequencies) according to physical distance between variants, in *C. grandiflora* and *D. melanogaster*. In color: LD is polarized based on the minor allele frequency (MAF) and averaged over pairs of sites at similar distances, for different MAF thresholds; In grey: LD is computed between neutral derived variants irrespective of their frequency and averaged over pairs of sites at similar distances.

**Figure S13:**
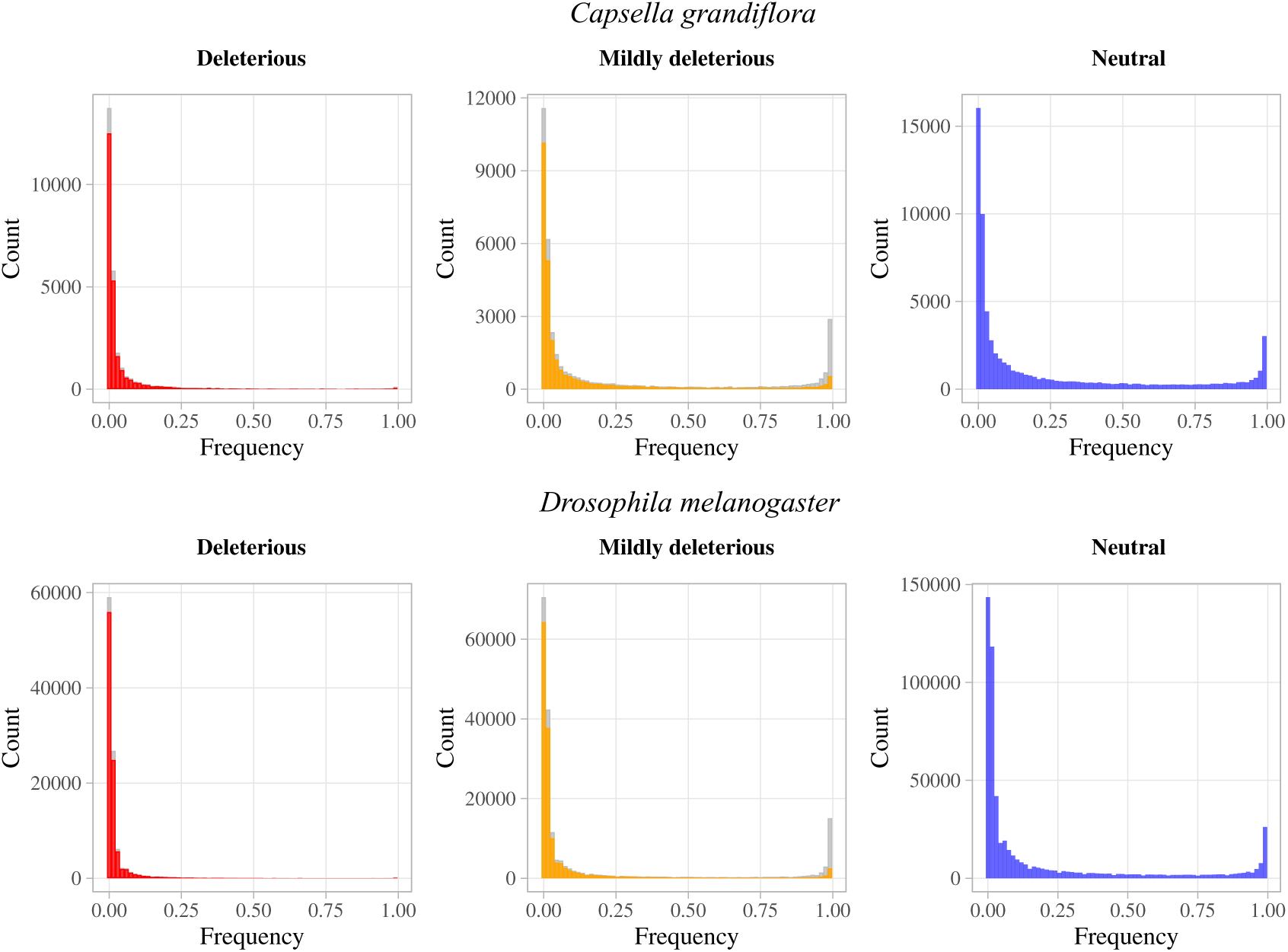
Site frequency spectrum of deleterious, mildly deleterious and neutral variants in *C. grandiflora* and *D. melanogaster* polarized by their derived state (in color, foreground) or by their SIFT score in the case of deleterious and mildly deleterious variants (in gray, background). The derived state of alleles was determined using the sequence of an outgroup (*Neslia paniculata* for *C. grandiflora* or *D. yakuba* for *D. melanogaster*, see Methods section). Polarizing by the SIFT score only slightly affected the SFS of deleterious alleles (as most derived mutations are also deleterious) and increased the number of high-frequency mildly deleterious mutations. Indeed, a proportion of high-frequency mildly deleterious mutations are inferred as ancestral, either because their are actually ancestral to the species and its outgroup or because they appeared in the lineage of the outgroup after its split with the lineage of *C. grandiflora*.

**Figure S14:**
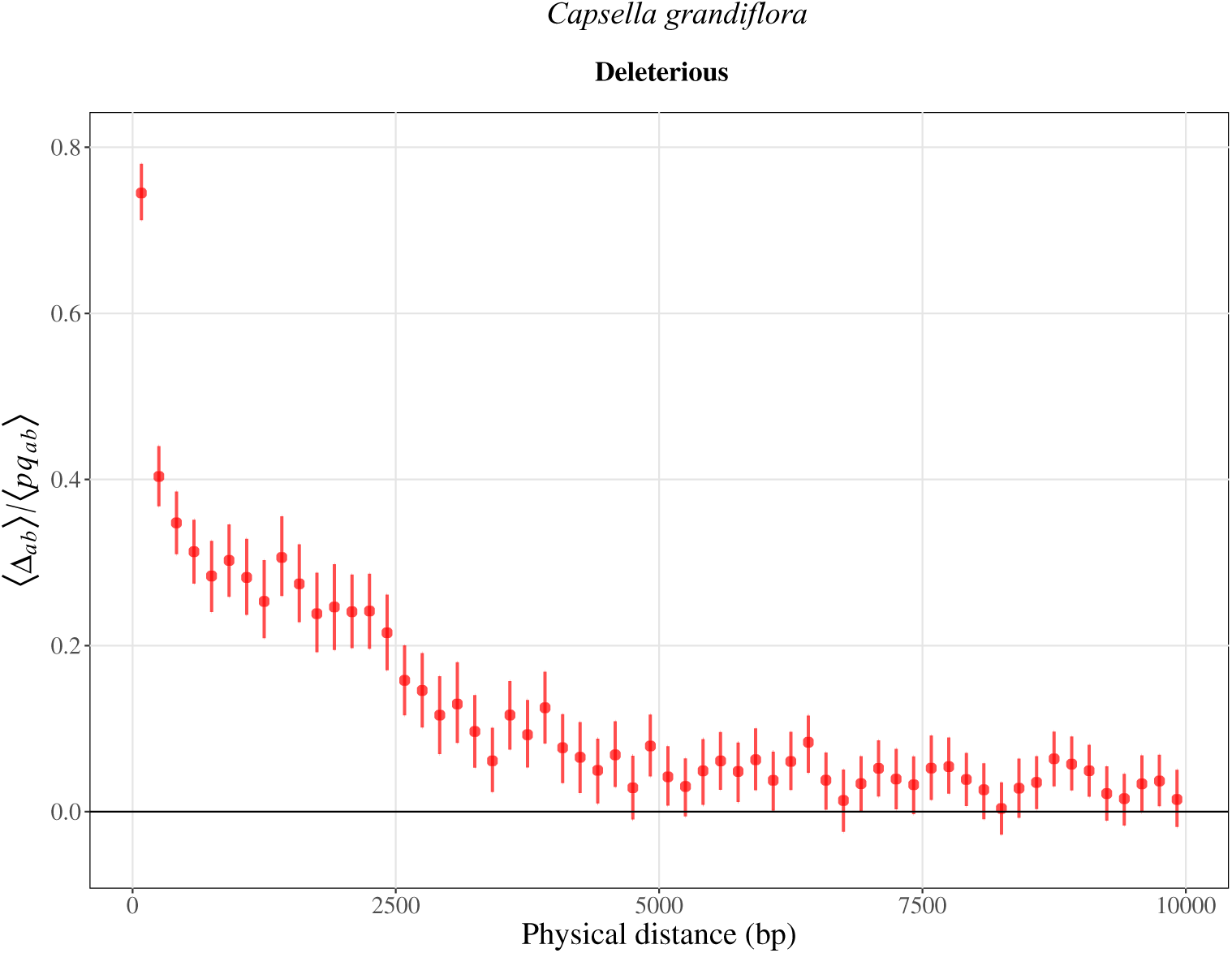
Average composite linkage disequilibrium Δ*_ab_* (scaled by the average product of allele frequencies) between deleterious variants according to physical distance in *C. grandiflora*. This plot corresponds to the top left panel in Figure 3 but LD is computed between pairs of SNPs up to 10kb and each distance classe is 167bp wide instead of 25bp wide in Figure 3.

**Figure S15:**
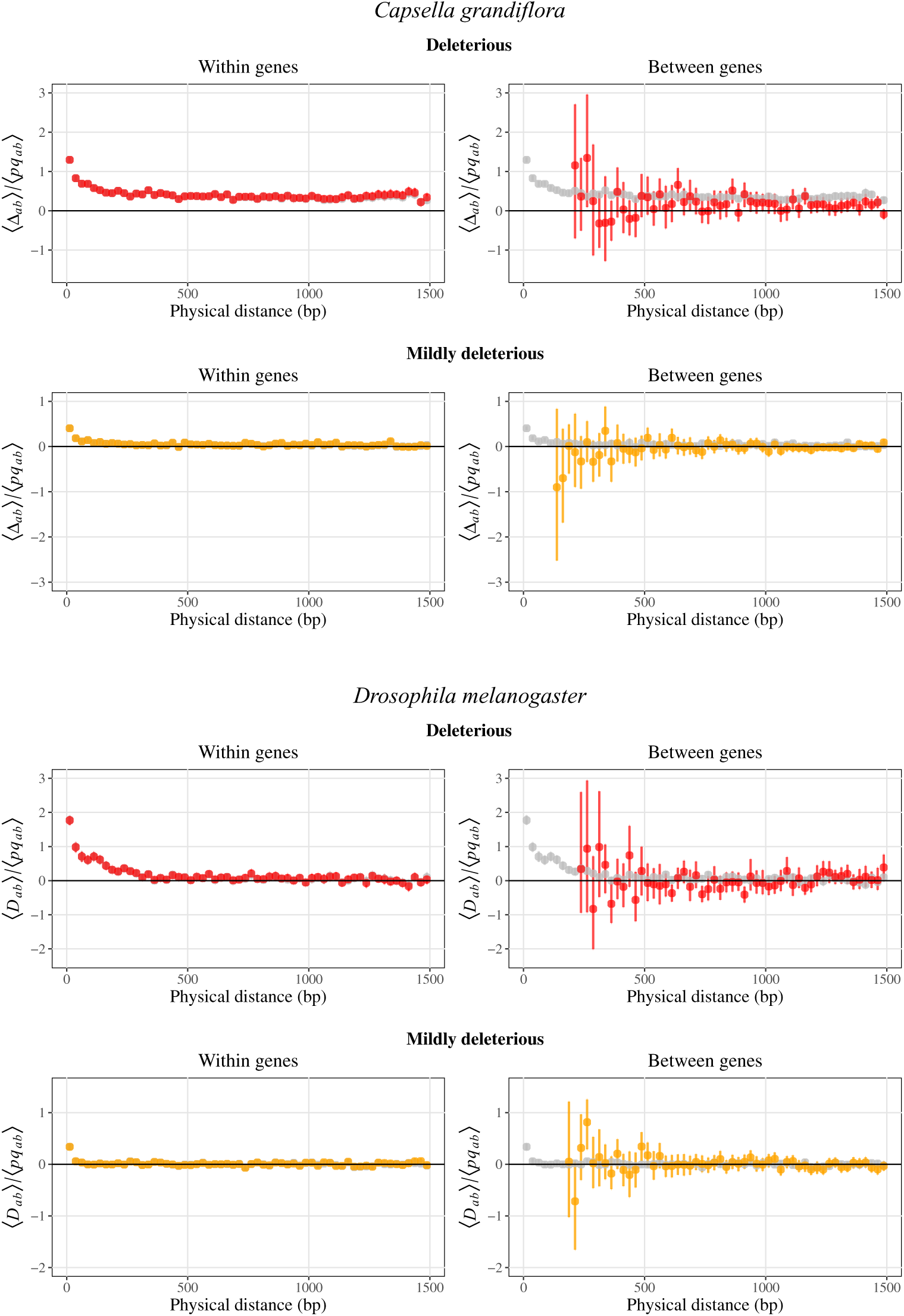
Average composite linkage disequilibrium Δ*_ab_* or linkage disequilibrium *D_ab_* (scaled by the average product of allele frequencies) between deleterious or mildly deleterious variants according to physical distance between variants, for pairs of mutations within or between genes in *C. grandiflora* and *D. melanogaster*. Points for distance classes with less than 20 pairs of mutations are not represented. Grey points are the same as in Figure 3 and are confounded with colored points in the ‘Within genes’ plots.

**Figure S16:**
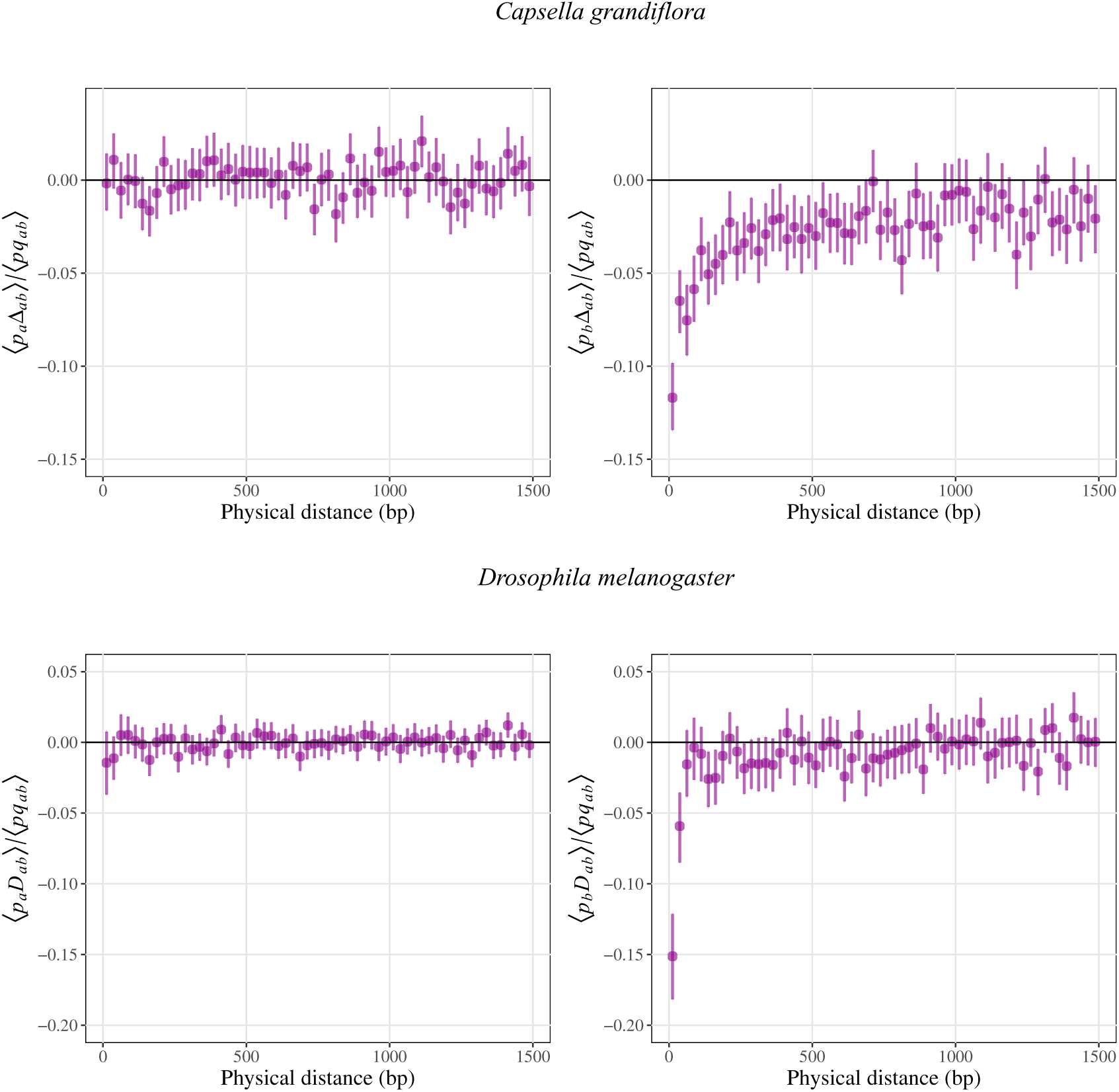
Averages of *p_a_*Δ*_ab_* (left panels) and *p_b_*Δ*_ab_* (right panels) (scaled by the average product of allele frequencies) over all pairs of deleterious alleles *a* and derived neutral alleles *b* at a given physical distance in *C. grandiflora* (top panels) and *D. melanogaster* (bottom panels).

**Figure S17:**
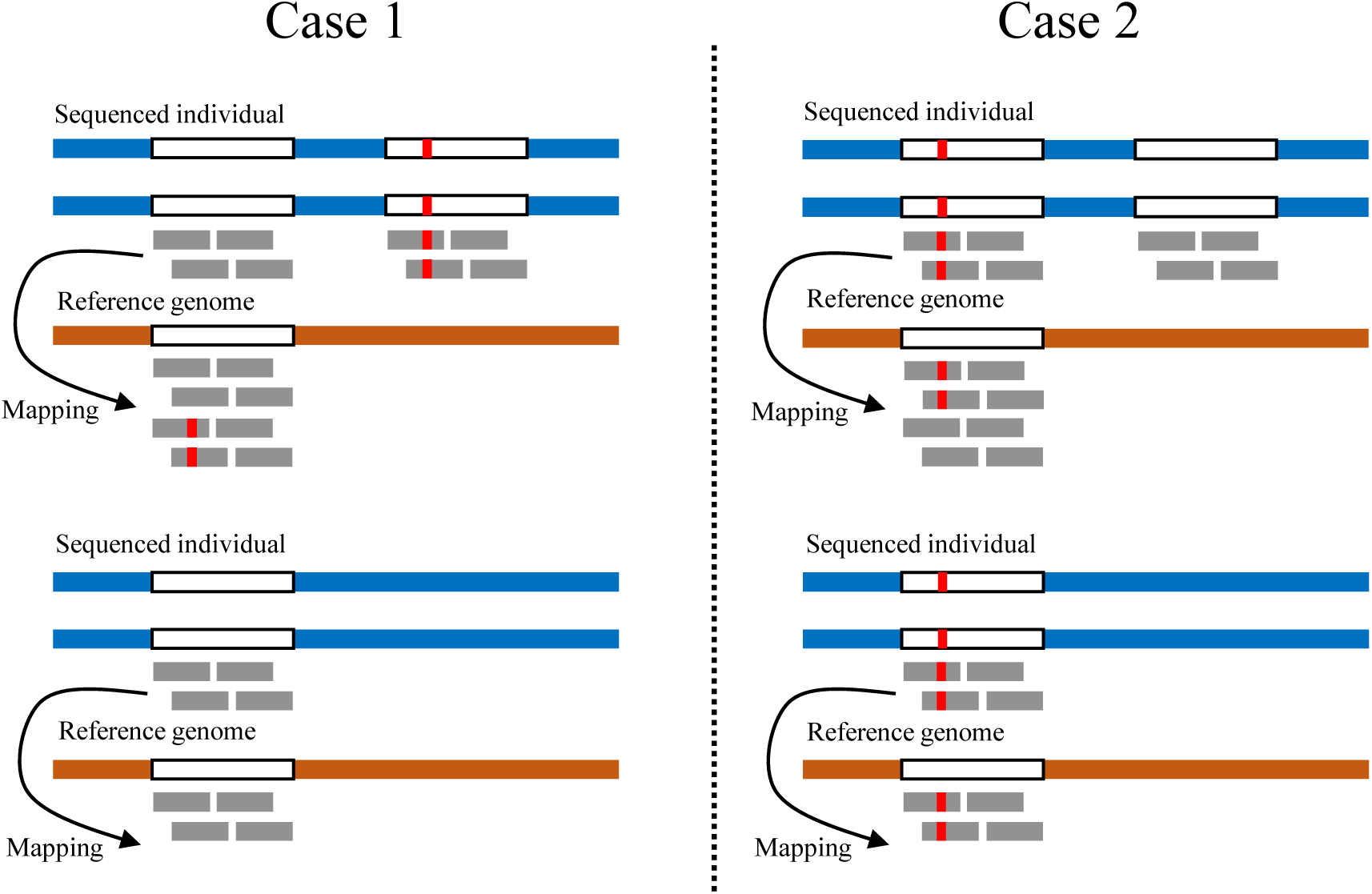
Cartoon illustrating the different genotypes found when mapping short reads from individuals carrying or not a duplication which is absent from the reference genome. To simplify the cartoon individuals are assumed to have homozygous genomes. In Case 1 a mutation is fixed in the duplicated region so that all individuals carrying the duplication appear as heterozygotes after mapping against the reference genome while individuals not carrying it appear as homozygous for the ancestral allele. In Case 2 a mutation (not present in the reference genome) is fixed in the original sequence but not the duplicated one, so that all individuals carrying the duplication appear as heterozygotes after mapping against the reference genome while individuals not carrying it appear as homozygous for the derived allele.

**Figure S18:**
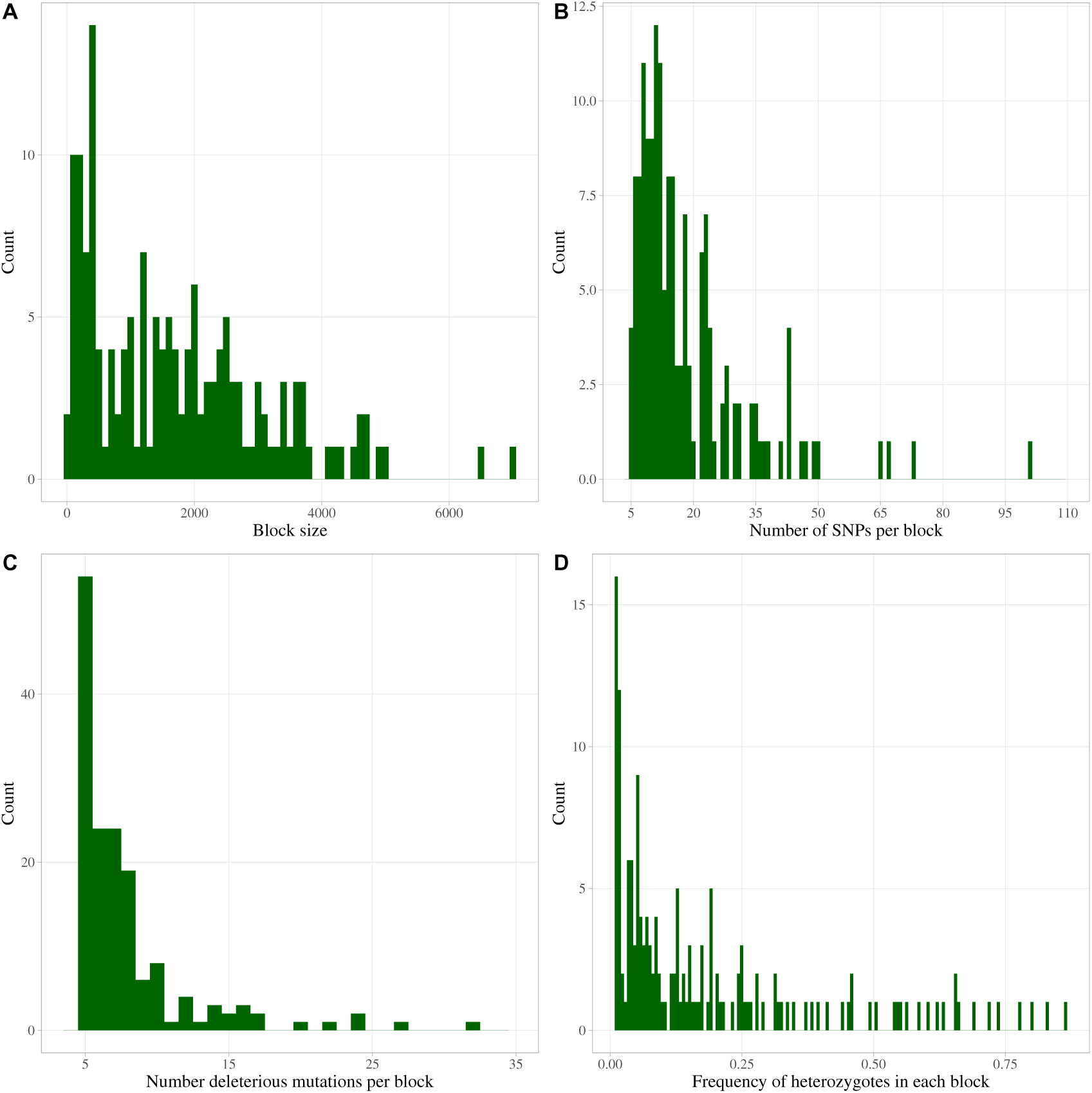
Distribution of various statistics related to heterozygosity blocks in the genome of *C. grandiflora*. A. Distribution of block size. B. Distribution of the number of SNPs per block. C. Distribution of the number of SNPs with deleterious mutations per block. D. Frequency spectrum of heterozygous individuals (*i.e.*, individuals potentially carrying the duplication) for each block.

**Figure S19:**
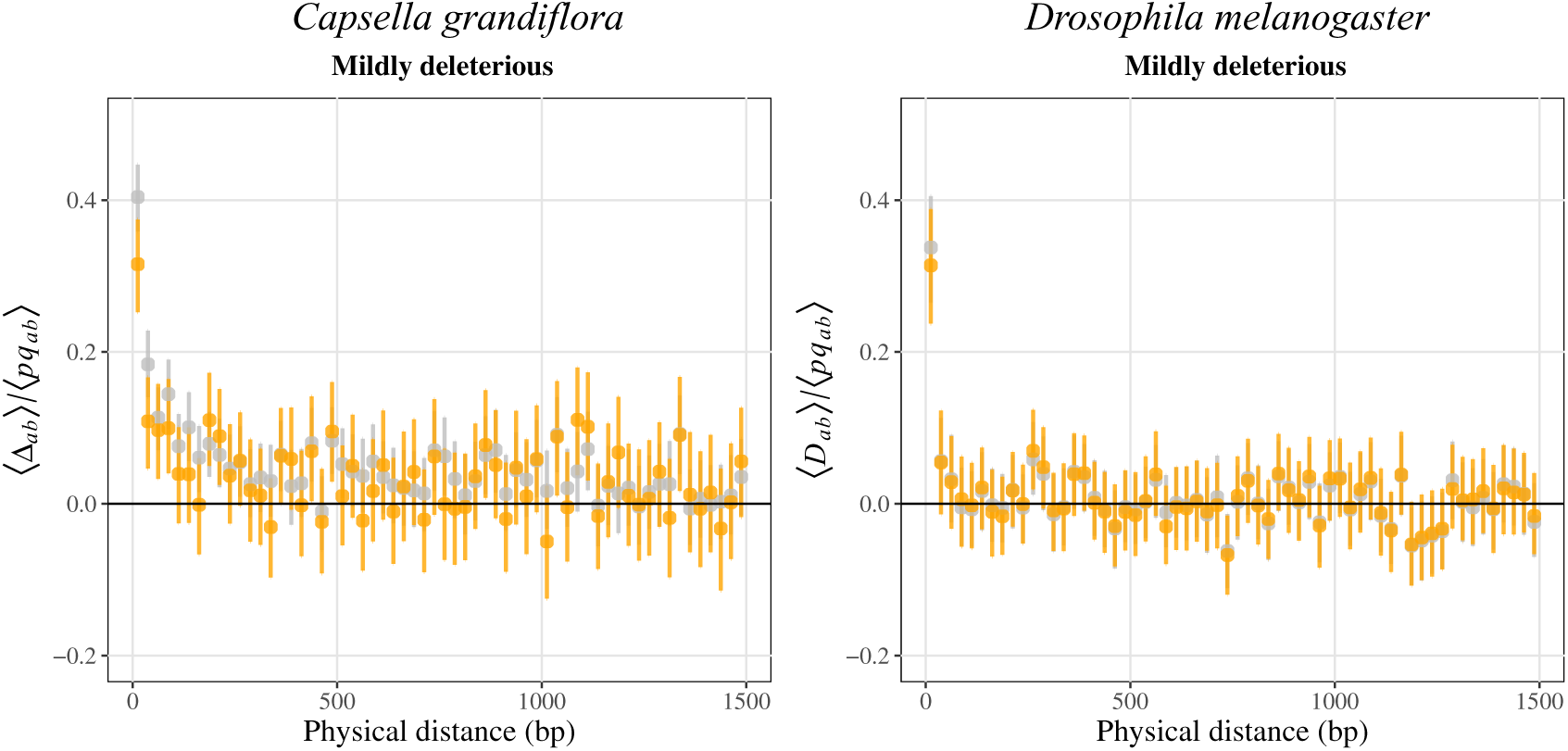
Left panel: Composite linkage disequilibrium Δ*_ab_* scaled by the product of allele frequencies between mildly deleterious variants according to physical distance between sites, after removing SNPs located in heterozygosity blocks (coloured) or considering all SNPs (grey, as in Figure 3) in *C. grandiflora*. Right pan rious varia chromosom

**Figure S20:**
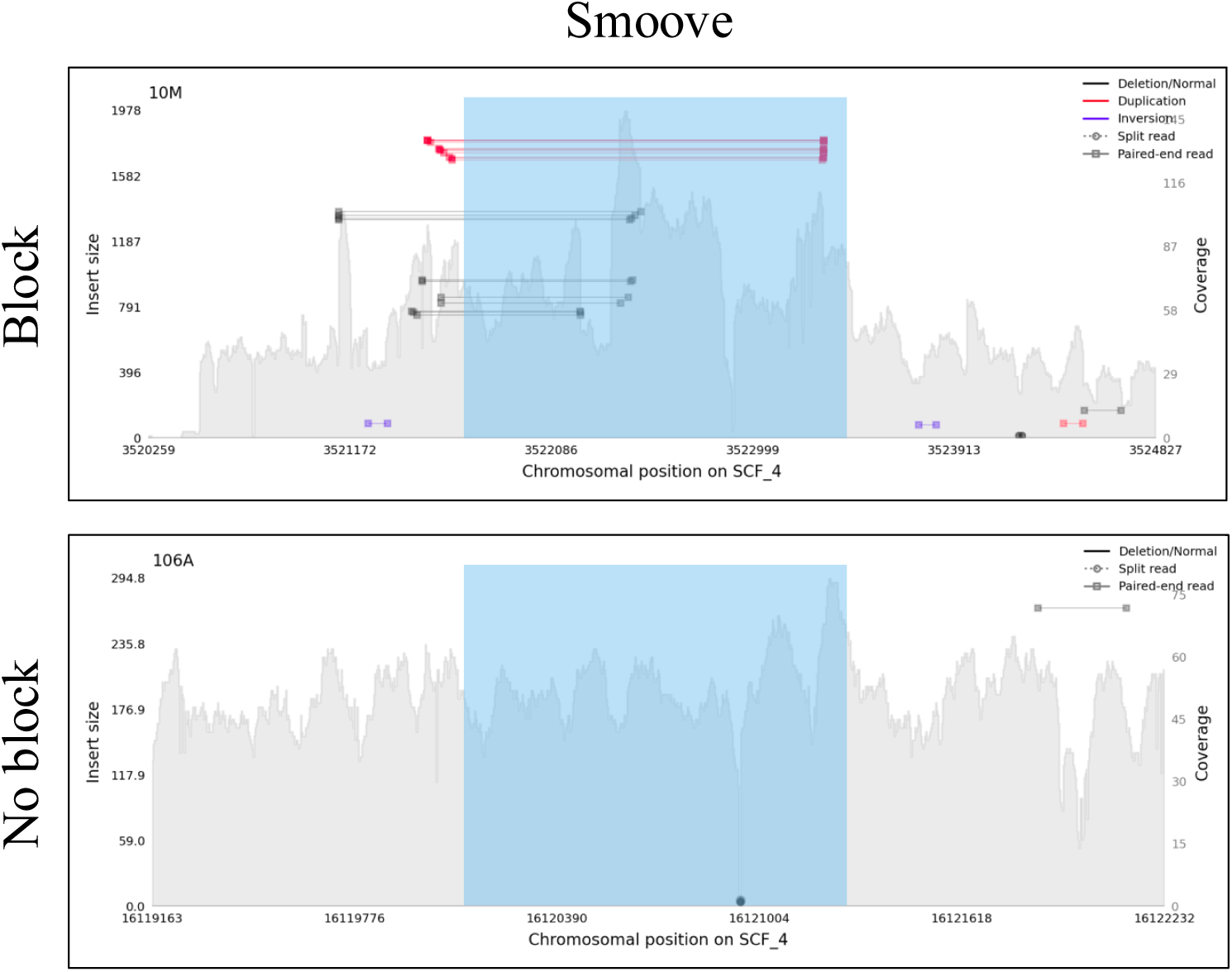
Example of a heterozygosity block (blue rectangle) located on chromosome 4 matching with a duplication inferred by Smoove and showing an elevated coverage in heterozygous individuals (10M; top panel) compared to homozygous individuals (106A; bottom panel).

**Figure S21:**
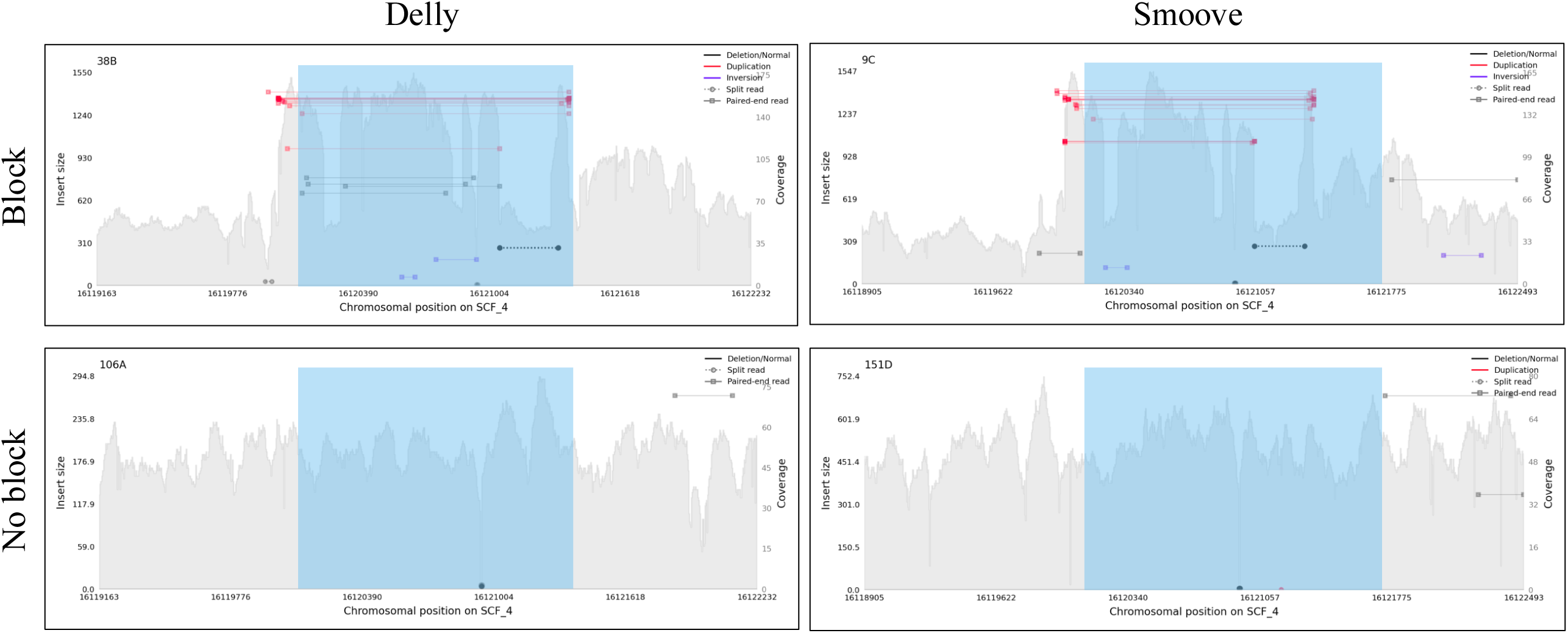
Example of a heterozygosity block (black horizontal bar) located on chromosome 4 matching with a duplication inferred by Delly (left panels) and Smoove (right panels) and showing an elevated coverage in heterozygous individuals (38B and 9C; top panels) compared to homozygous individuals (106A and 151D; bottom panels).

**Figure S22:**
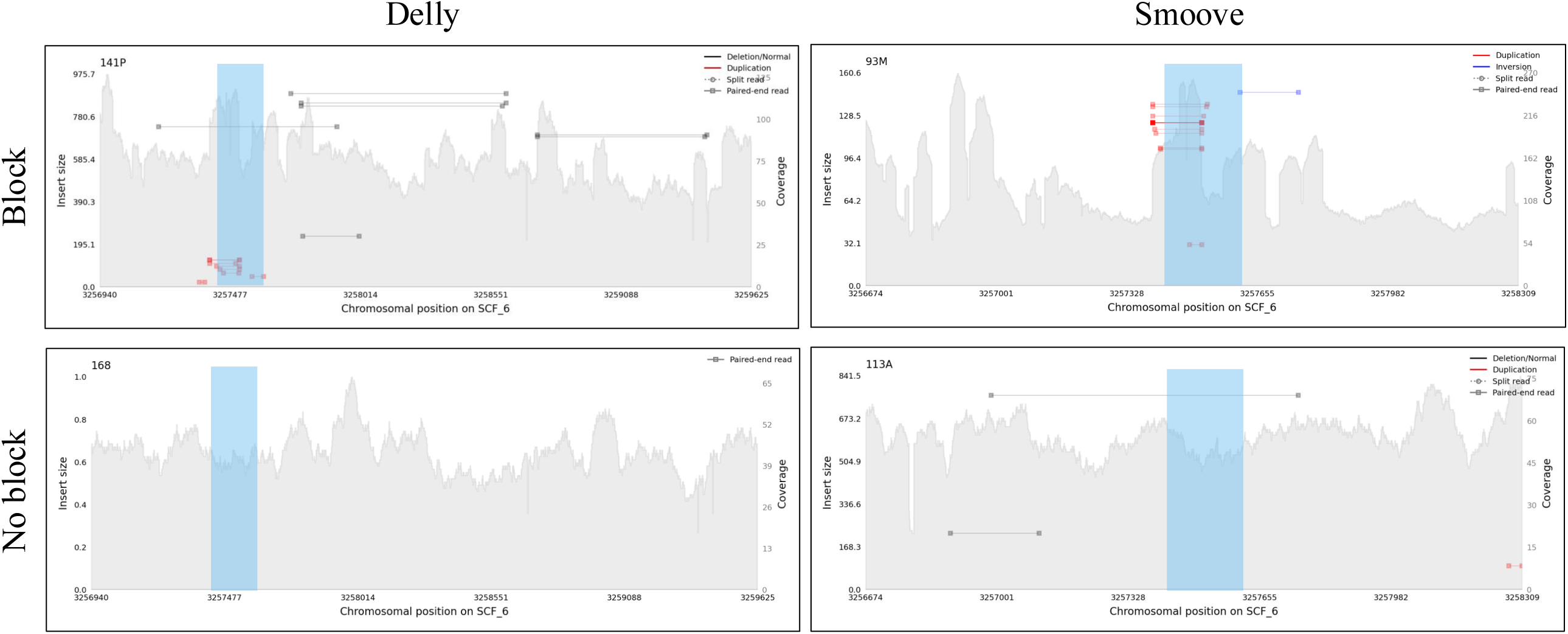
Example of a heterozygosity block (blue rectangle) located on chromosome 6 matching with a duplication inferred by Delly (left panels) and Smoove (right panels) and showing an elevated coverage in heterozygous individuals (141P and 93M; top panels) compared to homozygous individuals (168 and 113A; bottom panels).

**Figure S23:**
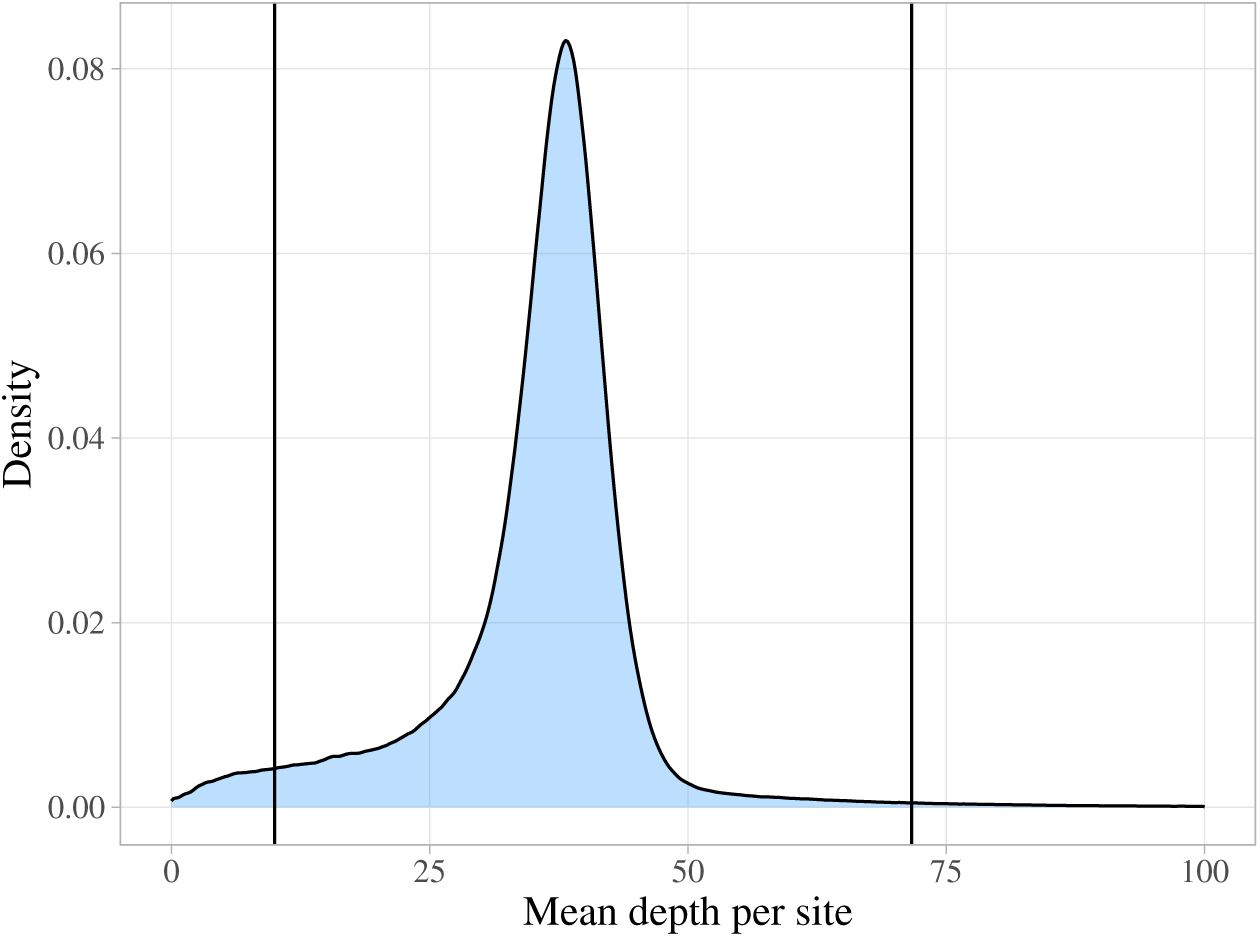
Distribution of mean depth per site and filtration criteria applied (vertical bars) on SNPs in *C. grandiflora*.

**Figure S24:**
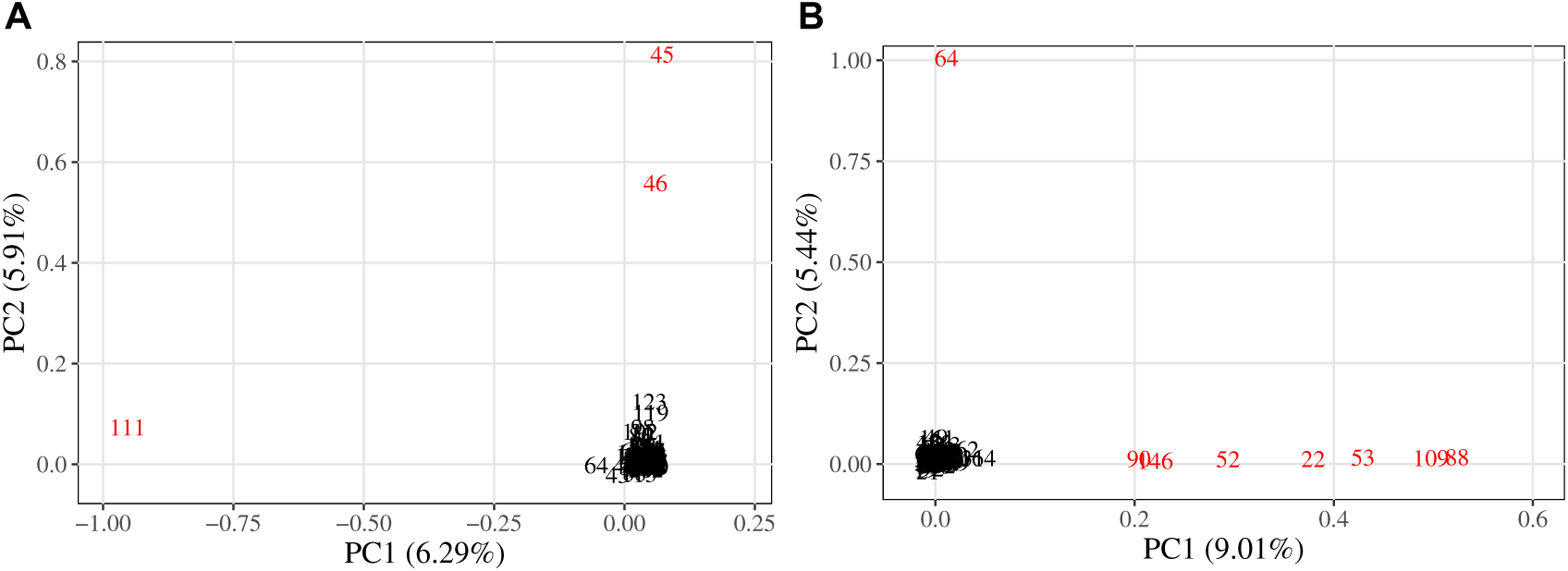
The two first components of the PCA on filtered SNPs of *C. grandiflora*. Red individuals in plot A where removed to draw plot B. The correspondence between individuals’ numbers and accession IDs is shown in Table S1. All red individuals correspond to diverged individuals that were excluded from LD computing. PCA was performed using Plink (Purcell et al., 2007) on independent SNPs extracted with the function *indep-pairwise* (window size = 50 kb, step size = 100 variants and *r*^2^ threshold = 0.1).

**Figure S25:**
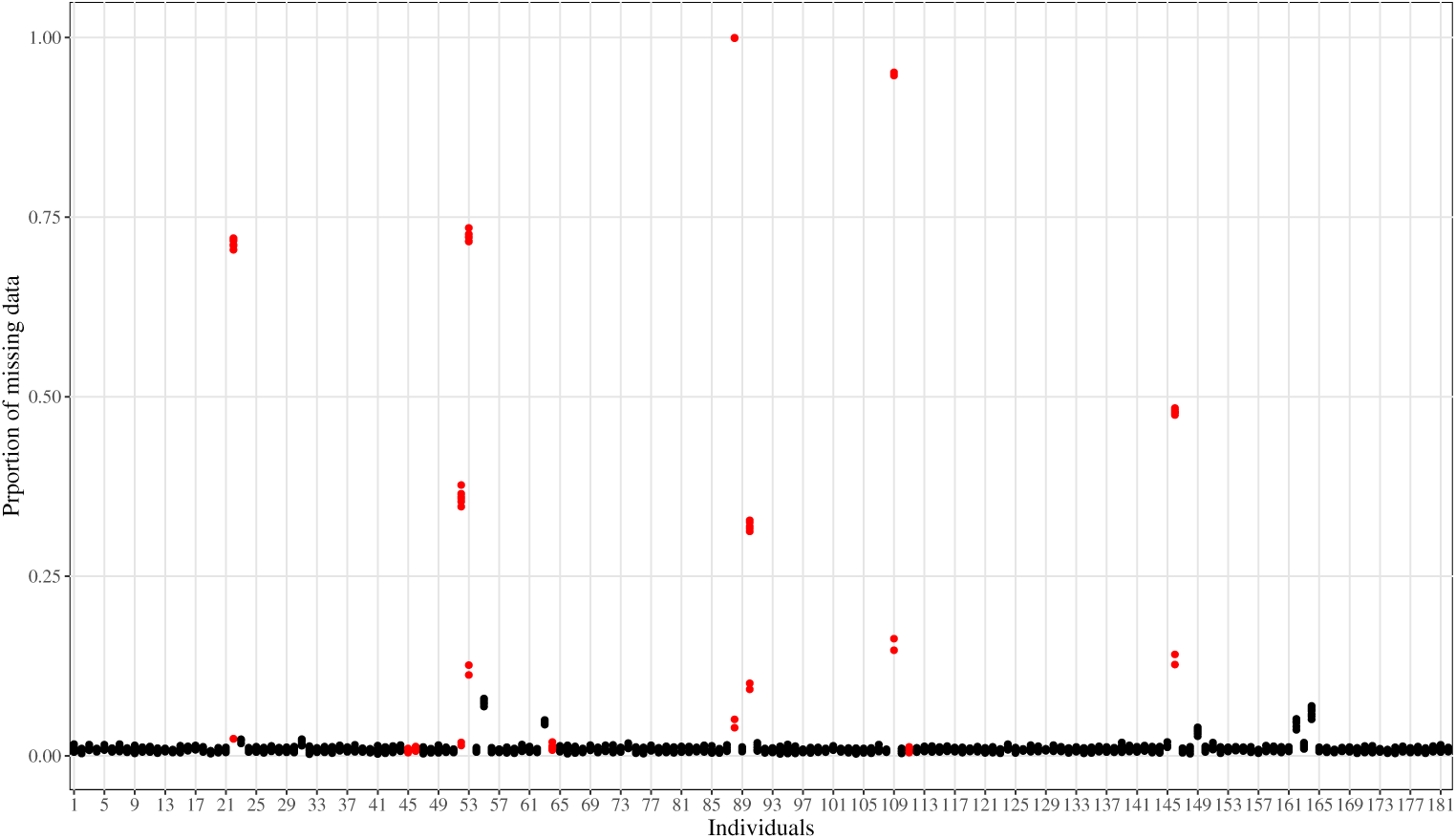
Proportion of sites with missing data in SNPs of *C. grandiflora* individuals. The correspondence between individuals’ numbers and accession IDs is shown in Table S1. Each point represents the proportion computed in each chromosome of each individual. Red points correspond to diverged individuals that were excluded from LD computing.

## Supplementary tables

**Table S1:** List of accessions of *Capsella grandiflora* with their ID from Josephs et al. (2015) SRA code, number used in the current paper, number of mapped reads and mean coverage after mapping and mean coverage of SNPs after filtering. Individuals shaded in grey correspond to diverged individuals that were removed from LD computing.

| ID | SRA | Number | Number of mapped reads | Mean coverage after mapping (X) | Mean coverage of SNPs after filtering (X) |
| --- | --- | --- | --- | --- | --- |
| 100O | SRR2065171 | 1 | 67,778,253 | 39.6265 | 38.9136 |
| 101S | SRR2065172 | 2 | 71,652,964 | 43.9521 | 38.8951 |
| 102K | SRR2065173 | 3 | 55,040,504 | 33.4317 | 29.9501 |
| 103K | SRR2065174 | 4 | 71,096,939 | 43.0686 | 40.5397 |
| 105G | SRR2065175 | 5 | 40,515,048 | 24.4133 | 23.1359 |
| 106A | SRR2065176 | 6 | 74,503,090 | 43.7571 | 41.785 |
| 107B | SRR2065177 | 7 | 72,820,349 | 43.9293 | 41.7436 |
| 108B | SRR2065178<br>SRR2065179 | 8 | 67,768,983 | 39.0761 | 35.4101 |
| 109E | SRR2065180 | 9 | 67,546,014 | 40.9459 | 37.6479 |
| 10M | SRR2065181 | 10 | 64,989,506 | 37.0686 | 35.6434 |
| 110M | SRR2065182 | 11 | 52,643,683 | 30.5764 | 30.3862 |
| 111C | SRR2065183<br>SRR2065184 | 12 | 143,035,188 | 83.04 | 68.3479 |
| 112H | SRR2065185 | 13 | 71,583,533 | 43.2113 | 40.8899 |
| 113A | SRR2065186 | 14 | 80,500,398 | 48.9141 | 43.5456 |
| 115L | SRR2065188 | 15 | 57,505,563 | 34.4798 | 33.3936 |
| 116L | SRR2070888 | 16 | 50,457,336 | 29.4382 | 29.5714 |
| 117R | SRR2065189 | 17 | 42,772,970 | 25.6085 | 24.8085 |
| 118W | SRR2065190 | 18 | 74,411,924 | 44.1095 | 43.7323 |
| 119M | SRR2070889 | 19 | 76,444,706 | 46.4393 | 41.4909 |
| 11G | SRR2065191 | 20 | 74,806,894 | 45.0182 | 40.4033 |
| 120C | SRR2070890 | 21 | 56,412,844 | 33.8994 | 32.5833 |
| 121B | SRR2065192 | 22 | 47,003,390 | 27.9812 | 6.70313 |
| 122 | SRR2070891 | 23 | 43,741,144 | 25.9645 | 19.5828 |
| 123D | SRR2065193 | 24 | 67,396,367 | 40.0744 | 34.8186 |
| 124Y | SRR2065194<br>SRR2065195 | 25 | 63,814,643 | 36.8642 | 37.6905 |
| 125X | SRR2065196 | 26 | 59,939,496 | 34.8289 | 33.5542 |
| 126Z | SRR2065197 | 27 | 60,874,245 | 35.3933 | 35.679 |
| 128P | SRR2065198 | 28 | 50,330,242 | 29.8525 | 28.9122 |
| 129Y | SRR2065199 | 29 | 59,748,060 | 35.8645 | 35.1082 |
| 130 | SRR2070892 | 30 | 71,681,229 | 41.7704 | 40.5615 |
| 131X | SRR2065200 | 31 | 42,373,447 | 25.33 | 22.7073 |
| 132L | SRR2065201 | 32 | 70,204,351 | 42.5192 | 41.3182 |
| 133V | SRR2065202 | 33 | 82,680,837 | 47.9079 | 47.9725 |
| 135F | SRR2065203 | 34 | 68,212,063 | 41.3666 | 39.9496 |
| 136R | SRR2065204 | 35 | 56,751,296 | 34.4575 | 34.9401 |
| 137Q | SRR2065205 | 36 | 54,998,156 | 31.8002 | 29.5052 |
| 138Q | SRR2065206 | 37 | 58,822,419 | 35.5121 | 28.9459 |
| 139G | SRR2065207 | 38 | 50,305,503 | 29.3703 | 29.2626 |
| 13L | SRR2065208 | 39 | 66,729,805 | 40.3389 | 41.1133 |
| 140S | SRR2065209 | 40 | 76,818,928 | 46.5473 | 47.3456 |
| 141P | SRR2065210 | 41 | 83,122,625 | 50.5074 | 49.5207 |
| 142B | SRR2065211 | 42 | 59,492,174 | 36.0179 | 34.982 |
| 143Y | SRR2065212 | 43 | 79,023,415 | 47.3287 | 43.0101 |
| 144Q | SRR2065213 | 44 | 59,298,700 | 34.2662 | 33.1329 |
| 145A | SRR2070893 | 45 | 65,754,737 | 38.299 | 35.6665 |
| 146J | SRR2065214 | 46 | 49,366,218 | 28.6793 | 25.9689 |
| 147Y | SRR2065215 | 47 | 69,417,464 | 42.0787 | 42.2985 |
| 148Z | SRR2065216 | 48 | 67,968,689 | 41.2659 | 41.5584 |
| 149 | SRR2070894 | 49 | 73,208,253 | 42.3707 | 40.701 |
| 14R | SRR2065217 | 50 | 63,299,238 | 36.9454 | 35.5008 |
| 151D | SRR2065218 | 51 | 77,462,657 | 46.6692 | 47.6571 |
| 152P | SRR2065219 | 52 | 55,738,604 | 33.0613 | 9.32303 |
| 153G | SRR2065220 | 53 | 42,178,619 | 25.0774 | 5.67162 |
| 154P | SRR2065221 | 54 | 68,051,515 | 40.8469 | 40.9878 |
| 155T | SRR2065222 | 55 | 22,682,911 | 13.5619 | 13.016 |
| 156R | SRR2065223 | 56 | 80,085,198 | 47.2397 | 48.0434 |
| 157x | SRR2065224 | 57 | 83,113,379 | 49.9481 | 52.9682 |
| 158J | SRR2065225 | 58 | 88,667,257 | 52.3153 | 50.3465 |
| 15U | SRR2065226 | 59 | 65,930,250 | 40.0612 | 37.384 |
| 160R | SRR2065227 | 60 | 53,656,225 | 31.8291 | 28.2538 |
| 161A | SRR2065228 | 61 | 81,200,362 | 48.6511 | 46.0047 |
| 162R | SRR2065229 | 62 | 76,273,152 | 45.8311 | 43.5795 |
| 163J | SRR2065230 | 63 | 41,555,559 | 24.857 | 11.8835 |
|  | SRR2065231 |  |  |  |  |
| 165I | SRR2065232 | 64 | 86,240,512 | 49.9489 | 25.7551 |
|  | SRR2065233 |  |  |  |  |
| 166 | SRR2070895 | 65 | 77,213,914 | 45.9567 | 44.6707 |
| 167Q | SRR2065234 | 66 | 55,064,212 | 32.8007 | 32.9024 |
| 168 | SRR2070896 | 67 | 64,985,455 | 38.7091 | 37.5054 |
| 16A | SRR2065235 | 68 | 128,526,954 | 75.3131 | 34.4697 |
|  | SRR2065236 |  |  |  |  |
| 170V | SRR2065237 | 69 | 101,435,487 | 58.9531 | 28.5943 |
|  | SRR2065238 |  |  |  |  |
| 172 | SRR2070897 | 70 | 83,336,374 | 50.4784 | 47.1442 |
| 173Z | SRR2065239 | 71 | 130,863,213 | 76.0802 | 36.1616 |
|  | SRR2065240 |  |  |  |  |
| 174I | SRR2065241 | 72 | 86,513,634 | 50.3249 | 47.6696 |
| 175D | SRR2065242 | 73 | 107,408,534 | 64.3758 | 32.1062 |
|  | SRR2065243 |  |  |  |  |
| 176X | SRR2065244 | 74 | 42,929,153 | 25.6369 | 21.7436 |
| 177K | SRR2065245 | 75 | 69,027,380 | 41.2525 | 40.3573 |
| 178M | SRR2065246 | 76 | 92,120,628 | 55.7699 | 54.9876 |
| 179E | SRR2070898 | 77 | 45,270,540 | 27.4644 | 25.3947 |
| 17Q | SRR2065247 | 78 | 79,781,641 | 47.1432 | 48.2977 |
| 180T | SRR2070899 | 79 | 135,713,967 | 81.2489 | 36.0529 |
|  | SRR2070900 |  |  |  |  |
| 181Y | SRR2065248 | 80 | 68,588,317 | 40.8242 | 39.8429 |
| 182Q | SRR2065249 | 81 | 65,624,998 | 37.856 | 36.7498 |
|  | SRR2065250 |  |  |  |  |
| 183E | SRR2065251 | 82 | 45,199,313 | 26.4439 | 27.0142 |
| 184M | SRR2065252 | 83 | 64,417,831 | 37.2486 | 36.6804 |
|  | SRR2065253 |  |  |  |  |
| 186W | SRR2065254 | 84 | 82,085,974 | 49.7373 | 52.0428 |
| 187K | SRR2065255 | 85 | 57,803,629 | 34.4179 | 28.187 |
| 189B | SRR2070901 | 86 | 60,620,390 | 35.7028 | 34.3665 |
| 18C | SRR2065256 | 87 | 70,251,238 | 41.5032 | 40.9202 |
| 190U | SRR2065257 | 88 | 45,087,744 | 26.9096 | 4.76871 |
| 192H | SRR2065258 | 89 | 53,042,382 | 31.2101 | 30.3332 |
| 193N | SRR2065259 | 90 | 40,187,845 | 23.9157 | 10.3354 |
| 195L | SRR2065260 | 91 | 38,309,552 | 22.344 | 22.5537 |
| 196X | SRR2070903 | 92 | 45,285,316 | 26.469 | 27.0444 |
| 197L | SRR2065261 | 93 | 83,198,562 | 47.3261 | 47.3051 |
| 198A | SRR2065262 | 94 | 76,537,570 | 45.1522 | 44.7034 |
| 199 | SRR2070904 | 95 | 64,112,895 | 37.8811 | 36.712 |
| 19Y | SRR2065263 | 96 | 63,152,265 | 38.2398 | 40.4914 |
| 1N | SRR2065264 | 97 | 123,532,118 | 72.4085 | 36.0577 |
|  | SRR2065265 |  |  |  |  |
| 2 | SRR2070905 | 98 | 79,710,259 | 46.6854 | 45.512 |
| 200U | SRR2065266 | 99 | 69,154,039 | 40.9279 | 39.8612 |
| 202I | SRR2065267 | 100 | 63,451,416 | 38.1595 | 35.4487 |
| 203A | SRR2065268 | 101 | 46,773,953 | 28.3813 | 25.1764 |
| 204O | SRR2065269 | 102 | 122,273,541 | 74.2437 | 62.6971 |
| 207D | SRR2070906 | 103 | 64,959,938 | 37.8733 | 36.9108 |
| 208 | SRR2070907 | 104 | 70,009,972 | 41.3305 | 41.5862 |
| 209P | SRR2070908 | 105 | 81,473,140 | 49.291 | 51.0841 |
| 20B | SRR2065270 | 106 | 73,916,835 | 44.7042 | 44.918 |
| 23Z | SRR2065271 | 107 | 58,846,003 | 34.3074 | 34.7468 |
| 24F | SRR2065272 | 108 | 70,901,904 | 42.1701 | 40.0409 |
| 25I | SRR2065273 | 109 | 30,966,308 | 18.4625 | 3.72872 |
| 26F | SRR2065274 | 110 | 81,127,053 | 49.0621 | 49.8611 |
| 27K | SRR2065275 | 111 | 70,841,232 | 41.9656 | 38.0147 |
| 28C | SRR2065276 | 112 | 67,652,969 | 41.0223 | 38.2379 |
| 29A | SRR2065277 | 113 | 52,360,150 | 30.168 | 31.2883 |
| 3 | SRR2070909 | 114 | 70,868,057 | 42.4081 | 40.2711 |
| 30Y | SRR2065278 | 115 | 57,257,392 | 33.935 | 34.9868 |
| 31T | SRR2065279 | 116 | 66,956,787 | 40.5396 | 39.6854 |
| 32 | SRR2070910 | 117 | 63,974,853 | 36.8332 | 36.3046 |
| 33E | SRR2065280 | 118 | 66,269,515 | 40.136 | 39.7893 |
| 34D | SRR2065281 | 119 | 65,381,523 | 38.8589 | 37.5032 |
| 35F | SRR2065282 | 120 | 78,697,770 | 46.7425 | 43.332 |
| 36H | SRR2065283 | 121 | 63,378,792 | 38.2639 | 39.0714 |
| 37 | SRR2070911 | 122 | 51,697,048 | 29.7453 | 29.7603 |
| 38B | SRR2065284 | 123 | 74,775,822 | 44.6129 | 46.4256 |
| 39N | SRR2065285 | 124 | 43,221,649 | 24.772 | 25.1637 |
| 41J | SRR2065286 | 125 | 73,532,236 | 43.6266 | 41.1315 |
| 42 | SRR2070912 | 126 | 64,804,156 | 39.1906 | 38.811 |
| 43F | SRR2065287 | 127 | 61,837,342 | 35.0002 | 28.9142 |
| 44W | SRR2065288 | 128 | 69,967,806 | 41.3651 | 23.2308 |
| 45 | SRR2070913 | 129 | 68,586,777 | 40.6128 | 37.9552 |
| 46 | SRR2070914 | 130 | 63,373,419 | 36.4663 | 35.6593 |
| 470x19 | SRR2070915 | 131 | 73,794,006 | 43.3242 | 28.2146 |
| 47K | SRR2065289 | 132 | 77,879,463 | 47.0856 | 45.219 |
| 48 | SRR2070916 | 133 | 61,847,174 | 35.8292 | 34.3142 |
| 49Z | SRR2065290 | 134 | 68,981,492 | 41.5254 | 38.9901 |
| 4S | SRR2065291<br>SRR2070917 | 135 | 86,021,503 | 48.9558 | 43.2052 |
| 50S | SRR2065292 | 136 | 53,003,268 | 31.8066 | 32.5276 |
| 51N | SRR2065293 | 137 | 62,167,457 | 35.9878 | 35.1954 |
| 52A | SRR2065294 | 138 | 69,974,028 | 41.4568 | 38.7089 |
| 53 | SRR2070918 | 139 | 58,899,793 | 34.0147 | 32.2693 |
| 54T | SRR2065295 | 140 | 48,983,666 | 28.1313 | 29.0827 |
| 55Y | SRR2065296 | 141 | 74,374,540 | 42.797 | 41.1833 |
| 58Y | SRR2065297 | 142 | 38,233,744 | 23.1927 | 23.8191 |
| 59N | SRR2070919 | 143 | 95,923,790 | 57.7721 | 50.6596 |
| 5S | SRR2065298 | 144 | 56,562,999 | 33.1306 | 31.7725 |
| 60T | SRR2070920 | 145 | 41,103,610 | 24.3865 | 22.4176 |
| 61G | SRR2065299 | 146 | 31,608,046 | 19.5132 | 7.15663 |
| 63D | SRR2065300 | 147 | 89,221,512 | 53.8929 | 52.5131 |
| 64E | SRR2065301<br>SRR2065302 | 148 | 114,132,152 | 68.2621 | 65.943 |
| 65T | SRR2065303 | 149 | 27,506,803 | 15.8318 | 15.8143 |
| 66X | SRR2065304 | 150 | 52,376,358 | 30.2621 | 30.2495 |
| 67R | SRR2065305 | 151 | 37,006,004 | 23.291 | 20.1181 |
| 6H | SRR2070921<br>SRR2070922 | 152 | 109,129,547 | 61.2037 | 56.0368 |
| 70V | SRR2065306 | 153 | 65,599,439 | 38.091 | 34.9359 |
| 71W | SRR2065307 | 154 | 58,970,708 | 34.4069 | 34.5394 |
| 72G | SRR2065308 | 155 | 71,155,909 | 43.0962 | 38.483 |
| 74M | SRR2065309 | 156 | 70,228,412 | 42.526 | 38.5986 |
| 75H | SRR2065310 | 157 | 56,901,796 | 34.2997 | 32.6267 |
| 76J | SRR2065311 | 158 | 52,906,046 | 31.3695 | 32.3992 |
| 78M | SRR2065312 | 159 | 65,199,534 | 37.8668 | 35.1942 |
| 79G | SRR2065313 | 160 | 72,256,279 | 41.602 | 36.2624 |
| 7K | SRR2065314<br>SRR2070923 | 161 | 61,484,995 | 34.8942 | 31.5093 |
| 80G | SRR2065315 | 162 | 24,004,415 | 13.576 | 13.5362 |
| 81J | SRR2065316 | 163 | 42,228,487 | 24.708 | 23.1726 |
| 82J | SRR2065317 | 164 | 23,140,063 | 13.1211 | 12.7753 |
| 83Z | SRR2065318 | 165 | 51,511,789 | 30.558 | 31.7037 |
| 85I | SRR2065319 | 166 | 79,359,602 | 47.8465 | 45.5222 |
| 86I | SRR2065320 | 167 | 73,299,422 | 44.4755 | 44.3322 |
| 88 | SRR2070924 | 168 | 68,610,724 | 40.2406 | 38.111 |
| 89L | SRR2065321 | 169 | 79,240,635 | 48.0663 | 47.7996 |
| 8U | SRR2065322 | 170 | 71,987,770 | 42.7442 | 40.3343 |
| 90E | SRR2065323 | 171 | 56,934,227 | 33.6398 | 33.7031 |
| 91C | SRR2065324 | 172 | 58,279,751 | 34.4491 | 34.4027 |
| 91x35 | SRR2070925 | 173 | 75,035,018 | 45.1561 | 46.6624 |
| 92H | SRR2065325 | 174 | 156,190,901 | 93.1882 | 86.4507 |
|  | SRR2065326 |  |  |  |  |
| 93M | SRR2065327 | 175 | 103,788,836 | 62.2141 | 64.942 |
| 94U | SRR2065328 | 176 | 52,791,072 | 31.3697 | 25.3255 |
| 95O | SRR2065329 | 177 | 76,074,792 | 46.1104 | 43.8904 |
| 96O | SRR2065330 | 178 | 70,086,066 | 42.3797 | 40.5477 |
| 97C | SRR2065331 | 179 | 100,347,463 | 57.7404 | 54.8625 |
|  | SRR2065332 |  |  |  |  |
| 98S | SRR2065333 | 180 | 55,438,898 | 32.4998 | 30.0239 |
| 99X | SRR2065334 | 181 | 48,525,235 | 28.3407 | 28.1156 |
| 9C | SRR2065335 | 182 | 70,004,164 | 41.2806 | 39.1199 |

**Table S2:** Structural variants called with Delly and Smoove that overlap with blocks of heterozygosity. Note that SVs can be of more than one type when Delly and Smoove detected different SV types.

| Scaffold number | Delly | Smoove | Both | Block number | Block start | Block end | Duplication | Deletion | Inversion | Unknown |
| --- | --- | --- | --- | --- | --- | --- | --- | --- | --- | --- |
| 1 | 1 | 0 | 0 | 51 | 243485 | 245445 | 0 | 0 | 1 | 0 |
| 1 | 0 | 0 | 1 | 126 | 336834 | 337847 | 0 | 1 | 0 | 1 |
| 1 | 1 | 0 | 0 | 322 | 603255 | 606258 | 0 | 0 | 1 | 0 |
| 1 | 0 | 0 | 1 | 549 | 847548 | 849702 | 0 | 1 | 0 | 0 |
| 1 | 1 | 0 | 0 | 898 | 1247070 | 1247243 | 0 | 0 | 1 | 0 |
| 1 | 0 | 0 | 1 | 1509 | 1957980 | 1959493 | 0 | 1 | 0 | 0 |
| 1 | 1 | 0 | 0 | 3383 | 3991057 | 3991412 | 0 | 0 | 0 | 1 |
| 1 | 0 | 0 | 1 | 6656 | 7539847 | 7543540 | 1 | 0 | 0 | 0 |
| 1 | 0 | 0 | 1 | 7253 | 8158786 | 8163531 | 0 | 1 | 1 | 0 |
| 1 | 1 | 0 | 0 | 7879 | 8773641 | 8776177 | 0 | 1 | 0 | 0 |
| 1 | 1 | 0 | 0 | 7896 | 8775752 | 8776184 | 0 | 0 | 1 | 0 |
| 1 | 0 | 1 | 0 | 8147 | 9012447 | 9017083 | 0 | 1 | 0 | 0 |
| 1 | 1 | 0 | 0 | 9765 | 10750119 | 10751167 | 0 | 1 | 0 | 0 |
| 1 | 1 | 0 | 0 | 11960 | 21886568 | 21889997 | 0 | 1 | 0 | 0 |
| 1 | 0 | 0 | 1 | 15161 | 27098904 | 27100568 | 0 | 1 | 0 | 0 |
| 2 | 0 | 0 | 1 | 4623 | 11903424 | 11909888 | 0 | 1 | 0 | 0 |
| 2 | 1 | 0 | 0 | 4649 | 11912346 | 11912595 | 0 | 1 | 0 | 0 |
| 3 | 0 | 0 | 1 | 1638 | 2360914 | 2363989 | 0 | 1 | 1 | 0 |
| 3 | 0 | 0 | 1 | 1889 | 2597139 | 2601720 | 0 | 1 | 0 | 0 |
| 3 | 0 | 0 | 1 | 5739 | 6783453 | 6786609 | 0 | 1 | 0 | 0 |
| 3 | 0 | 0 | 1 | 5820 | 6861615 | 6865684 | 0 | 1 | 0 | 0 |
| 3 | 1 | 0 | 0 | 6679 | 7675680 | 7676616 | 0 | 1 | 0 | 0 |
| 3 | 0 | 0 | 1 | 8088 | 9280004 | 9282621 | 1 | 0 | 0 | 0 |
| 3 | 0 | 0 | 1 | 8191 | 9402903 | 9406476 | 1 | 0 | 0 | 0 |
| 3 | 0 | 0 | 1 | 9164 | 12310950 | 12311825 | 0 | 1 | 0 | 1 |
| 3 | 1 | 0 | 0 | 11105 | 16156986 | 16158400 | 1 | 0 | 0 | 0 |
| 4 | 0 | 1 | 0 | 2034 | 3521661 | 3523402 | 1 | 0 | 0 | 0 |
| 4 | 1 | 0 | 0 | 7517 | 14282076 | 14285059 | 1 | 0 | 0 | 0 |
| 4 | 0 | 0 | 1 | 7907 | 14869630 | 14871575 | 0 | 1 | 0 | 0 |
| 4 | 0 | 0 | 1 | 8041 | 15004330 | 15006703 | 0 | 1 | 0 | 0 |
| 4 | 0 | 0 | 1 | 9030 | 16120116 | 16121750 | 1 | 0 | 0 | 0 |
| 5 | 1 | 0 | 0 | 5287 | 10421913 | 10425469 | 0 | 1 | 0 | 0 |
| 5 | 0 | 0 | 1 | 7080 | 12284036 | 12286639 | 1 | 0 | 0 | 0 |
| 5 | 1 | 0 | 0 | 7718 | 13045459 | 13047694 | 1 | 0 | 0 | 0 |
| 5 | 0 | 1 | 0 | 7119 | 12320140 | 12320224 | 0 | 0 | 0 | 1 |
| 5 | 0 | 1 | 0 | 7700 | 13035704 | 13038392 | 0 | 1 | 0 | 0 |
| 5 | 0 | 1 | 0 | 7718 | 13045459 | 13047694 | 1 | 0 | 0 | 0 |
| 5 | 0 | 0 | 1 | 8478 | 13728556 | 13730927 | 0 | 1 | 0 | 0 |
| 6 | 0 | 0 | 1 | 2544 | 3257426 | 3257618 | 1 | 0 | 0 | 0 |
| 6 | 0 | 1 | 0 | 5715 | 6760657 | 6763138 | 0 | 0 | 0 | 1 |
| 6 | 0 | 1 | 0 | 7861 | 9255955 | 9256836 | 0 | 1 | 0 | 0 |
| 6 | 0 | 0 | 1 | 9228 | 15906739 | 15907682 | 0 | 1 | 0 | 0 |
| 7 | 1 | 0 | 0 | 7033 | 7517598 | 7517976 | 1 | 0 | 0 | 0 |
| 7 | 1 | 0 | 0 | 7036 | 7517788 | 7517969 | 1 | 0 | 0 | 0 |
| 7 | 0 | 1 | 0 | 10094 | 9974254 | 9979280 | 1 | 0 | 0 | 0 |
| 7 | 1 | 0 | 0 | 10096 | 9974327 | 9977315 | 0 | 1 | 0 | 0 |
| 8 | 0 | 0 | 1 | 3222 | 9906198 | 9907207 | 1 | 0 | 0 | 0 |
| 8 | 1 | 0 | 0 | 4481 | 11374860 | 11376037 | 0 | 0 | 1 | 0 |
| 8 | 0 | 0 | 1 | 4586 | 11551052 | 11554723 | 1 | 0 | 1 | 0 |
| 8 | 1 | 0 | 0 | 8889 | 16240125 | 16242147 | 0 | 1 | 0 | 0 |
| 8 | 1 | 0 | 0 | 8892 | 16241640 | 16242119 | 0 | 1 | 0 | 0 |

